# PinkyCaMP a mScarlet-based calcium sensor with exceptional brightness, photostability, and multiplexing capabilities

**DOI:** 10.1101/2024.12.16.628673

**Authors:** Ryan Fink, Shosei Imai, Nala Gockel, German Lauer, Kim Renken, Jonas Wietek, Paul J. Lamothe-Molina, Falko Fuhrmann, Manuel Mittag, Tim Ziebarth, Annika Canziani, Martin Kubitschke, Vivien Kistmacher, Anny Kretschmer, Eva Sebastian, Dietmar Schmitz, Takuya Terai, Jan Gründemann, Sami Hassan, Tommaso Patriarchi, Andreas Reiner, Martin Fuhrmann, Robert E. Campbell, Olivia Andrea Masseck

## Abstract

Genetically encoded calcium (Ca^2+^) indicators (GECIs) are widely used for imaging neuronal activity, yet current limitations of existing red fluorescent GECIs have constrained their applicability. The inherently dim fluorescence and low signal-to-noise ratio of red-shifted GECIs have posed significant challenges. More critically, several red-fluorescent GECIs exhibit photoswitching when exposed to blue light, thereby limiting their applicability in all-optical experimental approaches. Here, we present the development of PinkyCaMP, the first mScarlet-based Ca^2+^ sensor that outperforms current red fluorescent sensors in brightness, photostability, signal-to-noise ratio, and compatibility with optogenetics and neurotransmitter imaging. PinkyCaMP is well-tolerated by neurons, showing no toxicity or aggregation, both *in vitro* and *in vivo*. All imaging approaches, including single-photon excitation methods such as fiber photometry, widefield imaging, miniscope imaging, as well as two-photon imaging in awake mice, are fully compatible with PinkyCaMP.

## Introduction

GECIs are widely used for *in vivo* imaging of neuronal populations^1^. Since the development of the first green fluorescent protein-based GECIs^2–4^, significant progress has been made in improving signal-to-noise ratios and kinetics, with advancements seen in GCaMP3 (ref. ^5^, GCaMP5 (ref.^6^), G-GECO1 (ref. ^7^), and GCaMP6 (ref.^8^). These improvements have culminated in the recent development of jGCaMP8, which offers ultra-fast kinetics and heightened sensitivity^9,10^. In contrast to green fluorescent GECIs, red-shifted sensors are generally preferable, as longer-wavelength light penetrates deeper into tissue and induces less phototoxicity. However, despite the advancements in green fluorescent biosensors, red-shifted GECIs still face limitations, such as lower brightness, reduced photostability, and photoswitching, as well as unique challenges like lysosomal accumulation, which have persisted since they were first reported (ref.^7^). Commonly used red GECIs, like jRCaMP1a,b^11^, R-GECO1 (ref.^7^), jRGECO1a^11^, XCaMP-R^12^, and RCaMP3 (ref.^13^), each have their own limitations.

All red fluorescent GECIs reported to date are derived from three distinct lineages of naturally occurring RFPs: R-GECO1 (ref.^7^) and its progeny are based on the *Discosoma sp*.-derived^14^ mApple^15^; jRCaMP1a,b^11^ are based on *Entacmaea quadricolor* eqFP611-derived^16^ mRuby^17^; and K-GECO1 and FR-GECO1 are based on *Entacmaea quadricolor* eqFP578-derived^18^ mKate^19^. While jRGECO1a, evolved from R-GECO1, is the brightest commonly used red GECI, it is still more than three times dimmer than GCaMP6s^11^ and although RCaMP3, the successor to jRGECO1a, is brighter than other red GECIs, it is still dim compared to green GECIs like GCaMP^13^. Based on circularly permuted mApple (cpmApple), jRGECO1a exhibits photoswitching under blue light, which complicates its use in combination with other tools, as blue-light excitation can falsely increase its brightness without reflecting actual Ca^2+^ changes. The same issue applies to the recently developed RCaMP3, which, despite being the brightest red GECI reported to date, is unsuitable for all-optical applications due to persistent photoswitching under blue light illumination. Additionally, red-shifted Ca^2+^ sensors based on R-GECO (mApple based), such as jRGECO1a, R-CaMP2 (ref.^20^), XCaMP-R^12^, and RCaMP3 (ref.^13^), inherit not only the photoswitching, but also all suffer from lysosomal accumulation^21^, limiting their use in combined imaging and optogenetic experiments.

Developing new red-shifted GECIs is therefore crucial to overcome these limitations. One promising candidate fluorescence protein for the development of red-shifted GECIs is mScarlet, known for its enhanced brightness and minimal photoswitching behavior^22^. First published in 2016, mScarlet quickly gained popularity as a red-fluorescent marker protein^22^. This protein, composed of 232 amino acids with a molecular weight of 26.4 kDa, is similar in size to other red fluorescent proteins^15,23^; however, it differs significantly in important structural properties such as its chromophore orientation^22^. mScarlet is one of the brightest known red-fluorescent proteins only outperformed recently by mScarlet3 (ref.^24^). mScarlet has a quantum yield of about 70%, while other monomeric red fluorescent proteins (RFPs) used in GECIs, such as mApple or mRuby have yields far below 50%^22^.

Additionally, mScarlet demonstrates negligible photoswitching behavior^22^.These attributes make mScarlet an excellent candidate for use in optogenetic tools and biosensors. However, possibly due to its limited structural similarity to other RFPs, no GECIs utilizing mScarlet have been developed yet.

Here we introduce PinkyCaMP, the first mScarlet-based Ca^2+^ indicator, offering improved signal-to-noise ratio, brightness, photostability, no photoswitching, and an exceptional change in absolute fluorescence upon Ca^2+^ binding. We demonstrate its compatibility with blue light optogenetics and simultaneous green fluorescence-based neuromodulator imaging through various *in vitro* and *in vivo* experiments. Additionally, we validate PinkyCaMP in two-photon imaging, highlighting its potential for advanced imaging applications.

## Results

### Development of an mScarlet-based GECIs

To develop an mScarlet-based GECI, we took inspiration from the design of previously reported GFP- and RFP-based GECIs. As a first step, we screened a large library of circularly permuted mScarlet (cpmScarlet) variants with different lengths and compositions of linkers connecting a calmodulin (CaM)-binding peptide (RS20 derived from R-GECO1)^7^ to the N-terminus and CaM (also derived from R-GECO1) to the C-terminus (**Figure 1a** and **Figure S1a**). Two promising prototypes were identified and designated as PinkyCaMP0.1a (brighter) and 0.1b (more responsive). To further improve the performance, the two prototypes were subjected to directed evolution **(Figure 1b** and **Figure S1b,c)**. After 12 rounds of library creation and screening in *E. coli*, which included assaying approximately 6,000 protein variants for brightness and response to Ca^2+^, we had three promising variants designated as PinkyCaMP0.9a (brighter), PinkyCaMP0.9b (more responsive), and PinkyCaMP0.9c (**Figure 1c**; a balance of brightness and responsiveness) (**Figure S2**).

**Figure 1:**
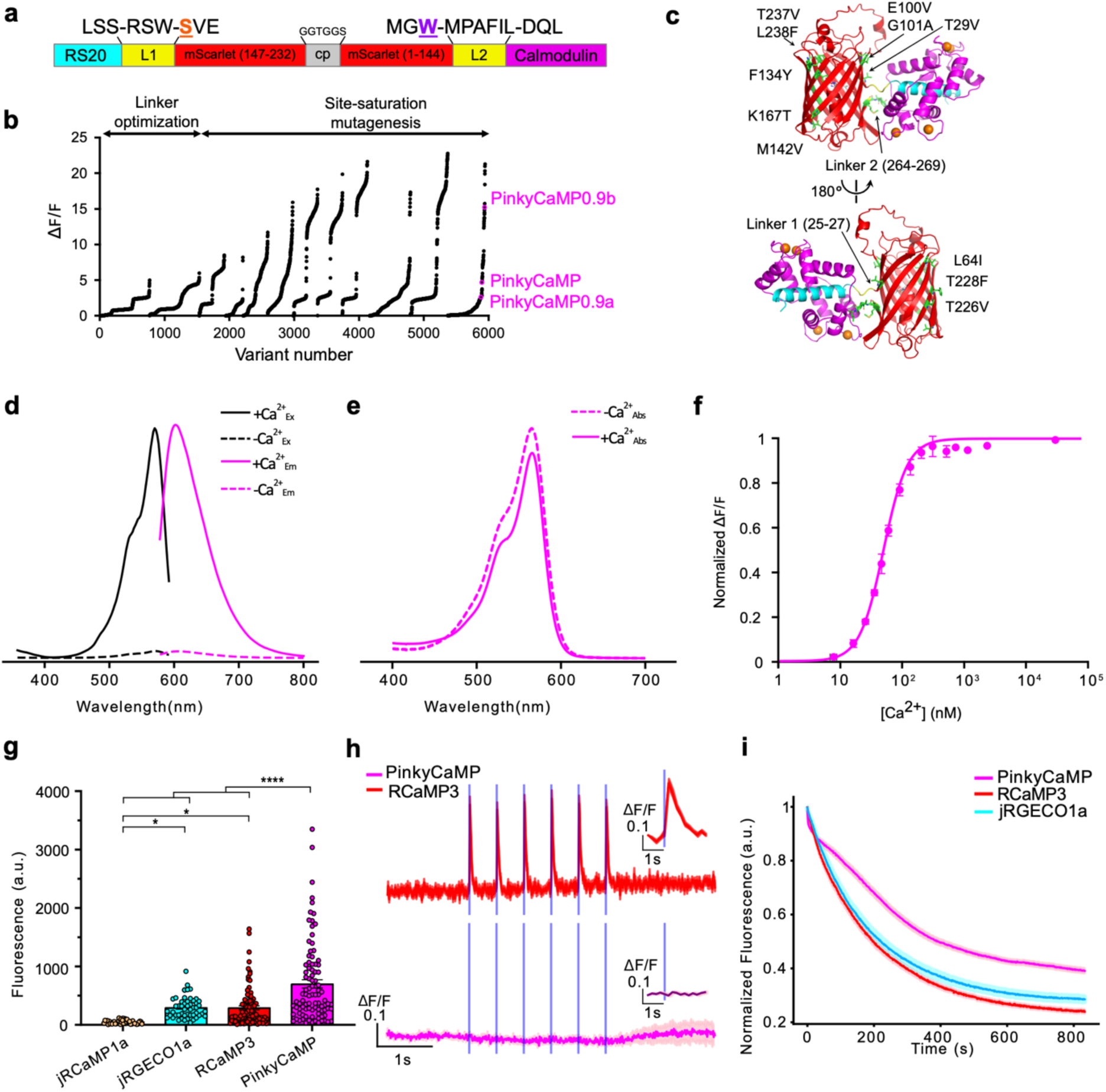
Development and characterization of PinkyCaMP. (a) Overview of the domain structure of PinkyCaMP. The two gate post residues<u>^62^</u> in cpmScarlet are shown in orange (S28) and purple (W144). (b) ΔF/F rank plot representing all crude proteins tested under the directed evolution screening conditions. ΔF/F values measured under these conditions are different from the values measured with purified proteins. (c) Modeled structure of PinkyCaMP with the positions of mutations indicated. RS20, cpmScarlet, CaM and linker residues are colored cyan, red, magenta and yellow respectively. The mutated residues are highlighted green. The model was prepared using AlphaFold3 (ref.<u>^63^</u>). (d) Excitation (emission at 620 nm) and emission (excitation at 540 nm) spectra of PinkyCaMP in the presence (39 µM) and absence of Ca2^+^. (e) Absorbance spectra of PinkyCaMP in the presence (39 µM) and absence of Ca2^+^. (f) Ca^2+^ titration curve of PinkyCaMP, n = 3 replicates (mean ± s.d.). (g) Baseline brightness in HEK cells expressing jRCaMP1a (50 ± 5 a.u.; n = 47 total cells), jRGECO1a (294 ± 25 a.u.; n = 50 total cells), RCaMP3 (292 ± 35 a.u.; n = 89 total cells), and PinkyCaMP (701 ± 71 a.u.; n = 93 total cells), all with three replicate measurements. One-way ANOVA with Tukey’s post-hoc test, **** p ≤ 0.001, * p ≤ 0.05 (mean ± s.e.m.). (h) Photoswitching of RCaMP3 (n = 33 cells) and PinkyCaMP (n = 20 cells), was assessed in HEK cells by imaging with constant 560 nm excitation light and periodic pulses of 470 nm light at 1 mW/mm2 and 0.1 Hz. Inset is an enlarged version of each first stimulation (mean ± s.e.m.). Three replicate measurements were performed for each sensor. (i) Averaged and normalized photostability traces of PinkyCaMP (n = 72 cells), RCaMP3 (n = 56 cells), and jRGECO1a (n = 76 total cells), from three replicate measurements (mean ± s.e.m).

### Characterization of mScarlet-based GECIs as purified proteins

To characterize the biophysical and photophysical properties of PinkyCaMP0.9a,b,c, we first expressed and purified each of the three proteins. Based on this characterization, as well as preliminary cell-based imaging studies, PinkyCaMP0.9c was selected as the best variant for its balance of brightness and responsiveness and renamed as PinkyCaMP (**Figure 1d,e,f** and the results of PinkyCaMP0.9a and 0.9b are shown in Figure S3). PinkyCaMP exhibits excitation and emission peaks of 568 and 600 nm, respectively, an absorbance peak of 567 nm, and a high ΔF/F of 15.1 (**Figure 1d,e** and **Table 1**). PinkyCaMP has a pKa of 6.83 and 4.24 with and without Ca^2+^ respectively (Figure S4 and Table 1), and an apparent Kd of 54 nM. These results indicate that, relative to other red GECIs, PinkyCaMP has the desirable characteristics of being less sensitive to cytoplasmic pH changes and having higher affinity for Ca2+ (Figure 1f, Figure S4a,d and Table 1). Upon binding to Ca2+, the quantum yield increases from 0.03 to 0.48, and the extinction coefficient decreases from 71,000 M-1cm-1 to 60,000 M^-1^cm^-1^ (**Table 1**). The intrinsic brightness of PinkyCaMP in the Ca^2+^-bound state matches that of jRCaMP1a which is the brightest, yet most poorly responsive, red GECI.

**Table 1.** Biophysical properties of purified proteins.

| Sensor | Condition | pK<sub>a</sub> | K<sub>d</sub><br>(nM) | Hill<br>coefficient | $\epsilon$<br>($\text{M}^{-1}\text{cm}^{-1}$) | $\phi$ | Brightness | $\Delta F/F^c$ |
| --- | --- | --- | --- | --- | --- | --- | --- | --- |
| PinkyCaMP | -Ca<sup>2+</sup> | 4.24 | 54 | 2.18 | 71,000 | 0.03 | 2.1 | 12.8 |
|  | +Ca<sup>2+</sup> | 6.83 |  |  | 60,000 | 0.48 | 28.9 |  |
| PinkyCaMP0.9a | -Ca<sup>2+</sup> | 5.66 | 149 | 1.30 | 89,000 | 0.16 | 14.3 | 2.0 |
|  | +Ca<sup>2+</sup> | 6.31 |  |  | 98,000 | 0.44 | 43.2 |  |
| PinkyCaMP0.9b | -Ca<sup>2+</sup> | 6.97 | 202 | 1.44 | 85,000 | 0.02 | 1.6 | 14.3 |
|  | +Ca<sup>2+</sup> | 6.64 |  |  | 57,000 | 0.43 | 24.4 |  |
| RCaMP3 | -Ca<sup>2+</sup> | 9.21 | 130 | 1.81 | 3,300 | 0.20 | 0.7 | 25.5 |
|  | +Ca<sup>2+</sup> | 5.79 |  |  | 56,400 | 0.31 | 17.5 |  |
| jRGECO1a<sup>a</sup> | -Ca<sup>2+</sup> | 8.6 | 148 | 1.90 | 6,180 | 0.18 | 1.1 | 9.2 |
|  | +Ca<sup>2+</sup> | 6.3 |  |  | 53,300 | 0.21 | 11.2 |  |
| jRCaMP1a<sup>a</sup> | -Ca<sup>2+</sup> | 5.6 | 214 | 0.86 | 33,800 | 0.29 | 9.8 | 2.0 |
|  | +Ca<sup>2+</sup> | 6.4 |  |  | 54,100 | 0.54 | 29.2 |  |

<sup>a</sup>Values are from Dana et al.<sup>11</sup> except for quantum yields which are from Molina et al.<sup>25</sup>.
<sup>b</sup>Brightness = $\epsilon \times \phi$
<sup>c</sup>Values calculated based on the brightnesses as listed in this table.

### Basic characterization of PinkyCaMP in cultured cells

To assess brightness of PinkyCaMP in mammalian cells, we expressed PinkyCaMP, jRCaMP1a, jRGECO1a, and RCaMP3 in HEK293T cells. As demonstrated for other GECIs, the inclusion of the bacterial expression plasmid-derived leader sequence (RSET) enhances GECI expression in mammalian cells^13^, and the inclusion of a nuclear export sequence (NES) prevents the mixing of slower or biphasic Ca^2+^ kinetics in the nucleus. Thus these two elements were incorporated into the final version of PinkyCaMP for expression and characterization in cultured cells, ex vivo, and in vivo. In HEK293T cells, PinkyCaMP showed a mean baseline fluorescence of 701 ± 71 a.u. and is more than twice as bright as RCaMP3 (292 ± 35 a.u.; p < 0.0001, one-way ANOVA) and jRGECO1a (294 ± 25 a.u.; p < 0.0001, one-way ANOVA) and fourteen times brighter than jRCaMP1a (50 ± 4 a.u.; p < 0.0001, one-way ANOVA; **Figure 1g**).

Given its mScarlet backbone, PinkyCaMP was hypothesized to have no photoswitching under blue light, which is a crucial feature for multiplexing imaging with other optogenetic tools. Surprisingly, the original description of RCaMP3 13 did not include data on photoswitching despite its jRGECO1a derived origins. To test this, we stimulated HEK293T cells expressing either sensor with 470 nm light (10 ms pulses, 1 mW/mm²). PinkyCaMP displayed no increase in ΔF/F, confirming an absence of photoswitching behavior. In contrast, RCaMP3 exhibited pronounced photoswitching, with ΔF/F increasing up to 0.43 ± 0.04 during stimulation, followed by a subsequent decay in fluorescence (**Figure 1h**, inset).

Lastly, we assessed the photostability of PinkyCaMP in comparison to other GECIs. HEK293T cells expressing the sensors were exposed to continuous 560 nm light (1 mW/mm²), and fluorescence decay was recorded and normalized to the peak fluorescence. PinkyCaMP retained nearly 40% of its initial fluorescence, outperforming RCaMP3 (23%) and jRGECO1a (28%) (**Figure 1i**). PinkyCaMP’s photobleaching half time (τ = 410.8 s) is more than double that of RCaMP3 (τ = 197.0 s) and much longer than jRGECO1a (τ = 222.3 s), highlighting its superior photostability under continuous illumination.

### Spectral multiplexing of PinkyCaMP and CoChR

Given PinkyCaMP’s useful biophysical properties, foremost the non-detectable blue light mediated photoswitching, we aimed to combine PinkyCaMP with a channelrhodopsin (ChR) in a single bicistronic construct (**Figure 2a**) to enable all-optical manipulation and readout of neuronal activity. PinkyCaMP was combined with the trafficking-enhanced and soma-targeted variant of the ChR CoChR^26,27^ (stCoChR = CoChR-TS-Kv2.1, **Figure 2a)** to allow for blue-light mediated photocurrent generation, while in parallel enabling imaging of Ca^2+^ activity using PinkyCaMP with orange light (**Figure 2b**). Low intensity activation of CoChR with 1 ms light pulses (**Figure 2b**, bottom) showed a maximum activity at 468 ± 2 nm (**Figure 2c**). In the following, we used narrow bandpass filtered blue light (438 ± 15 nm; to activate stCoChR) and orange light (586 ± 15 nm) for Ca^2+^-imaging via PinkyCaMP (**Figure 2c)** to enable multiplexing.

**Figure 2:**
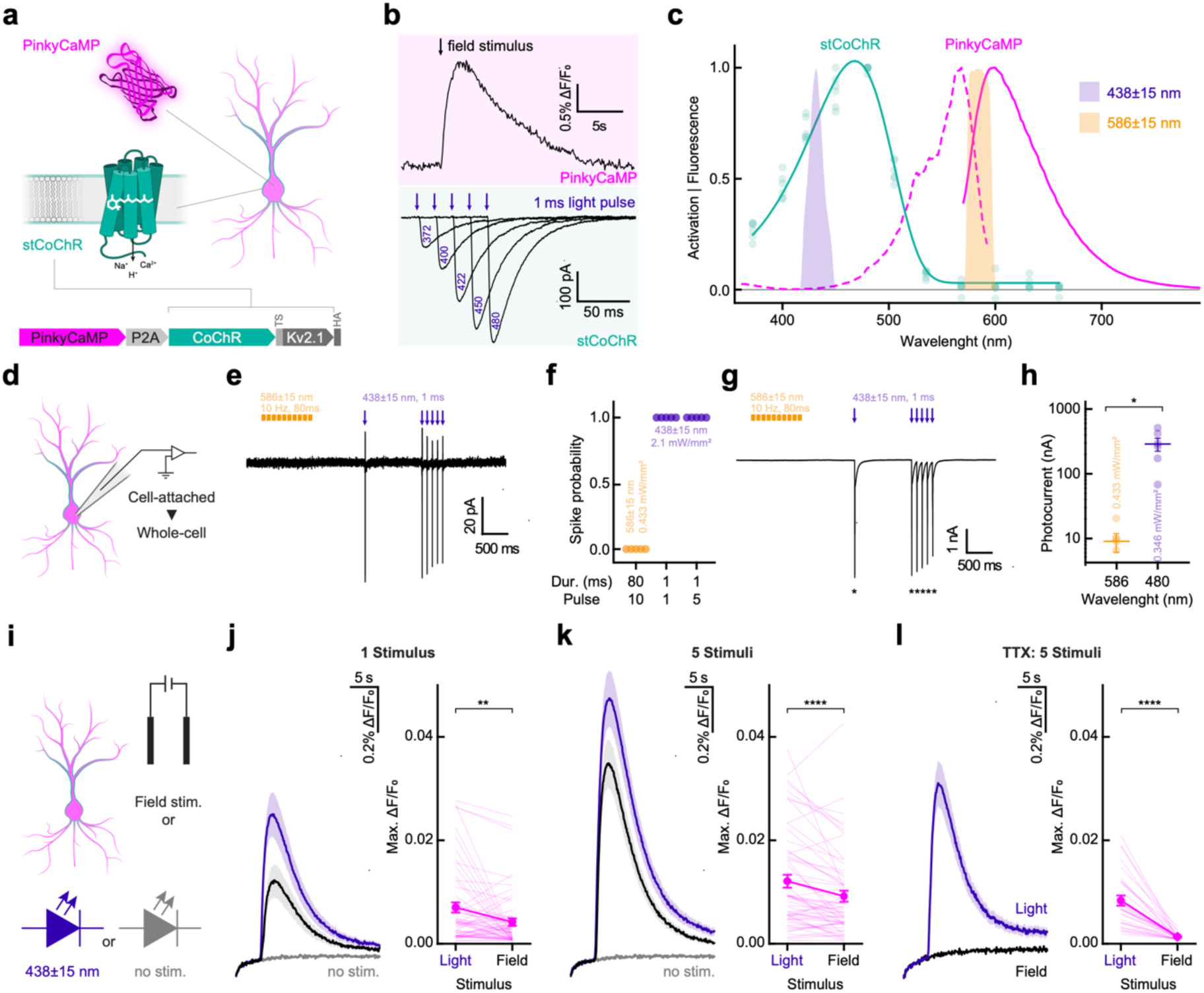
Spectral multiplexing of PinkyCaMP with CoChR. (a) Schematic of expressed proteins and neuronal localization (top) and construct design (bottom). (b) Single neuron Ca^2+^-imaging (top) with a single pulse field stimulation and electrophysiological current traces (bottom) elicited with different blue light applications. (c) Emission (solid, pink) and excitation (broken, pink) spectra of Ca^2+^-saturated PinkyCaMP together with the action spectrum recorded from PinkyCaMP-P2A-stCoCHR. n=6 cells. Stimulation light properties (blue and orange shaded areas) are shown. (d) Schematic of cell-attached / whole-cell measurements. (e) Representative cell-attached measurement. (f), Quantification of the spike probability in cell-attached mode under illumination conditions as shown in g. n=5 cells. (g) Representative whole-cell voltage-clamp measurement. * indicate action potentials. (h) Quantification of the generated photocurrent under imaging conditions in comparison to low intensity illumination used for action spectroscopy. Statistics: p=0.0312 n=6 cells. (i) Schematic of Ca^2+^ imaging with different stimuli: field stimulation (black), 438 nm LED (blue), and no stimulus (gray). (j) PinkyCaMP fluorescence change traces upon different single pulse stimuli (left) and quantification (right). Statistics: p=0.028, n=59 cells. Color coding as in i. (k) PinkyCaMP fluorescence change traces upon 5 consecutive stimuli (left) and quantification (right), same samples as in j. Statistics: p<0.0001, n=59. Color coding as in i. (l) PinkyCaMP fluorescence change traces upon 5 consecutive stimuli (left) and quantification (right) under 1 µM TTX treatment. n=30. Color coding as in i. All data are shown as mean ± SEM, and all statistical comparisons were performed as Wilcoxon matched-pairs signed rank tests.

Next, we assessed the electrophysiological impact caused by different light applications. We first measured action potential (AP) spiking activity in cell-attached configuration to maintain an unperturbed intracellular environment (**Figure 2d**). While imaging light of 586 nm did not cause any spiking, blue light at 438 nm caused reliable spiking when applied as a single pulse or at 10 Hz (**Figure 2e,f**). Following cell-attached measurements, we measured light-mediated photocurrents in whole-cell voltage-camp. Similarly to cell attached recording, blue light application caused AP firing(**Figure 2g**). However, the application of orange light used for imaging caused a small inward current of 9 ± 3 pA, much lower as currents measured for action spectra determination at 480 nm (288 ± 65 pA, **Figure 2h**).

In the next step we combined orange light PinkyCaMP imaging with stCoChR photoexcitation and additionally compared electric field stimuli on the same PinkyCaMP-2A-stCoChR expressing neurons (**Figure 2i**). A single 1 ms stimulus (either field stimulation or light application) caused a reliably detectable Ca^2+^ signal, while light evoked stimuli showed higher Ca^2+^ response on average (**Figure 2j**). The same neurons showed and increased Ca^2+^ response upon 5 delivered stimuli (at 10 Hz), while the tendency for a lower response (5 field stimuli vs. 5 light stimuli) persisted (**Figure 2k**). However, we consistently observed an up-ramping of the PinkyCaMP fluorescence signal during imaging, most noticeable at the start of the imaging session (**Figure 2j-l**). We anticipated that this small signal increase could be either caused by the small photocurrent generated by stCoChR (**Figure 2g**) and/or by the activation of voltage gated calcium channels and its inherent high Ca^2+^-permeability^27^ paired with the high sensitivity of PinkyCaMP (*K*_d_ = 54 nM). Indeed, when we blocked AP generation with TTX, no field stimulation-induced Ca^2+^-spikes were detectable anymore while light application still caused a significant Ca^2+^-signal (**Figure 2l**).

### Comparison of PinkyCaMP to other red GECIs

In order to compare PinkyCaMP to other red-fluorescent sensors and GCaMP6f under standardized conditions, we used mouse cortical slice cultures as an in situ model. In all cases, rAAV-mediated neuronal GECI expression resulted in readily identifiable Ca^2+^ transients, as shown for RCaMP3.0 and PinkyCaMP (Figure 3a,b; see Figure S5 for jRGECO1a, jRCaMP1a and GCaMP6f). These Ca2+ transients reflect spontaneous bursts of synchronized network activity, which are typical for slices of that age28,29. PinkyCaMP showed the highest baseline brightness (F0, **Figure 3c**) and large signal changes (ΔF/F0) (**Figure 3d**). In combination, these two properties give PinkyCaMP the highest absolute signal strengths of all tested red-fluorescent GECIs (**Figure 3e**), on par with GCaMP6f. Furthermore, PinkyCaMP showed relative long decay times (Figure 3f; τ = 4.9 ± 1.0 s, mean ± s.d.), similar to jRCaMP1a, and a high photostability (**Figure 3g**), whereas jRGECO1a and GCaMP6f showed significant bleaching under these conditions. Last, we compared how the different red-fluorescent GECIs were affected by additional blue-light (470 nm) illumination (**Figure S6** and **Figure 3h**). Here PinkyCaMP showed a stable signal after blue-light illumination, whereas jRGECO1a and RCaMP3.0 showed increased signals due to photoswitching which in amplitude were similar to synchronous events. The stability of PinkyCaMP was also confirmed in 60-min long-term recordings (Fig. S5 with comparisons).

**Figure 3:**
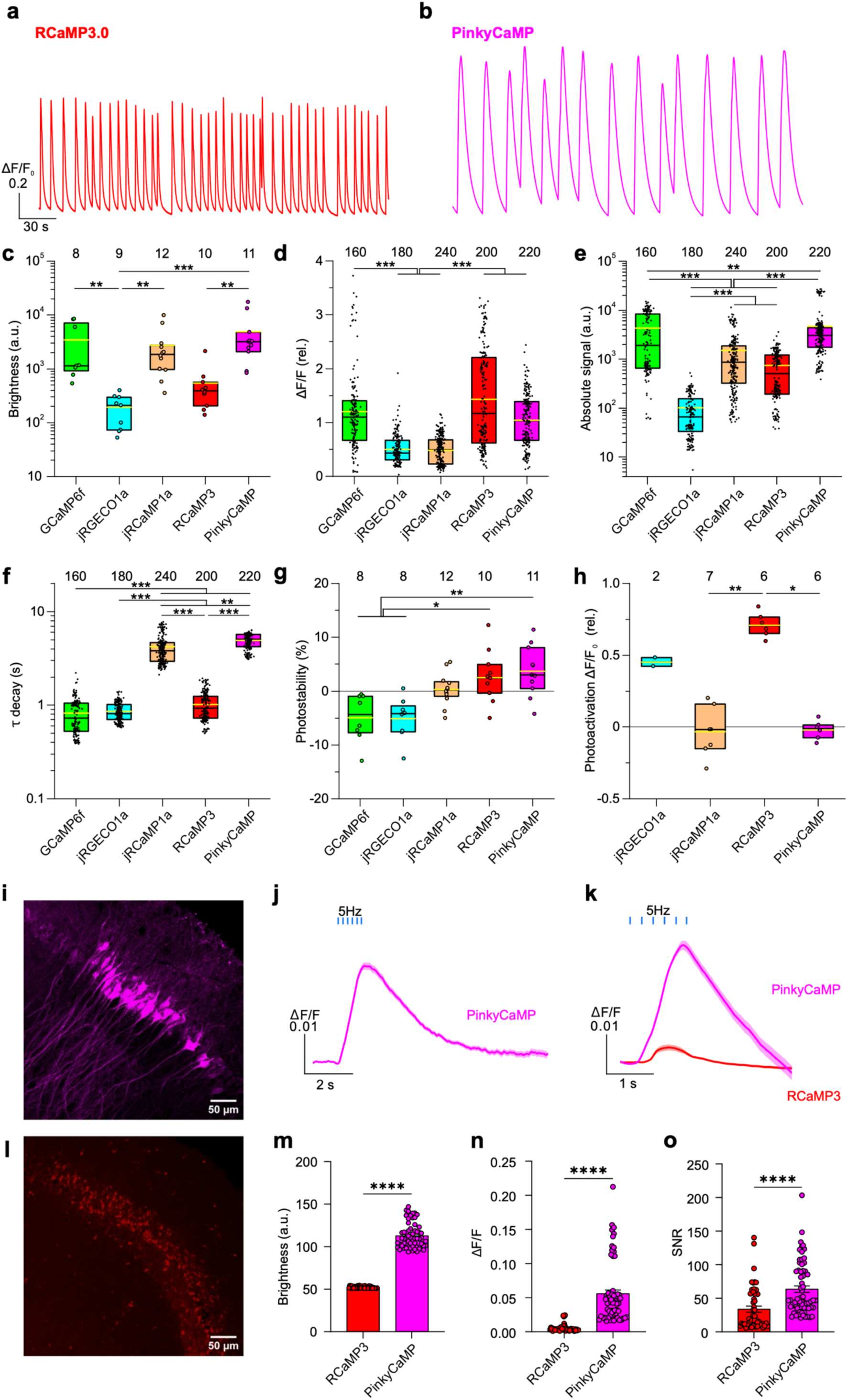
Comparison of PinkyCaMP and other GECIs in organotypic cultures and acute brain slices. (a,b) Spontaneous, synchronous calcium transients recorded with RCaMP3.0 and PinkyCaMP in cortical slice cultures from mouse (DIV 13-21; synapsin-driven expression after AAV-mediated transduction at DIV 1). (c) Fluorescence brightness (F0), (d) relative signal change (ΔF/F0), (e) absolute signal strength (F0aver·ΔF/F0), (f) decay time constants and (g) photostability for GCaMP6f, jRGECO1a, jRCaMP1a, RCaMP3.0 and PinkyCaMP. For other examples and parameters see Figure S5. (h) Assessment of blue-light photoswitching. For details see Figure S6. Black boxes indicate 25-75% percentiles and medians, the yellow lines means. The number of analyzed slices (c,g,h) or events (d,e,f) is given. Statistical differences are indicated with *p ≤ 0.05, **p ≤ 0.01, and ***p ≤0.001. (i) Confocal image of CA2 neurons in a 300 µm thick acute mouse brain slice expressing PinkyCaMP, scale bar 50 µm. (j) Average ΔF/F traces of 6 stimulation pulses at 5Hz stimulation for PinkyCaMP under low excitation light intensity (left; n=167 cells). (k) Average ΔF/F traces of 6 simulation pulses for PinkyCaMP and RCaMP3 at 5 Hz stimulation with high excitation light intensity (PinkyCaMP n=79 cells, RCaMP n=55 cells). (l) Confocal image of RCaMP3 in CA2 neurons, scale bar 50µm. (m) Baseline brightness of cells recorded with 10 Hz stimulation; PinkyCaMP: 106.250 ± 1.908, RCaMP3: 52.160 ± 0.171. (n) Maximal ΔF/F values for 10 Hz stimulation; PinkyCaMP: 0.0557 ± 0.0055, RCaMP3: 0.0051 ± 0.0007. (o) SNR at 10 Hz field stimulation; RCaMP3 33.711 ± 4.767, PinkyCaMP: 63.527 ± 7.880. (m-o) Values presented as mean ± s.e.m. PinkyCaMP n=60 cells, and RCaMP3 n=50 cells. Mann-Whitney U test, ****p ≤ 0.0001.

In addition, we performed a more detailed comparison between PinkyCaMP and RCaMP3 using widefield microscopy in acute hippocampal slices (**Figure S7a).** PinkyCaMP showed strong cytosolic and dendritic expression resulting in low-light recording capability (**Figure 3i,j**), with ΔF/F increasing significantly by field stimulation (**Figure S7b,c**). While neuropil fluorescence affected ΔF/F ratios, PinkyCaMP’s absolute fluorescent signal, photostability, and signal-to-noise ratio (SNR) remained high across trials. In contrast, RCaMP3 exhibited severe organellar, presumably lysosomal accumulation (**Figure 3l**) and required much higher light intensities (51 times higher than PinkyCaMP) for signal detection, therefore both sensors were measured with the higher light intensity (**Figure 3k** and **Figure S7d,e,f**). While PinkyCaMP demonstrated photobleaching from the high light intensity, it proved to be much more photostable than RCaMP3 as RCaMP3 failed to produce detectable signals after only a few replicate measurements. In contrast, cells expressing PinkyCaMP produced strong signals for dozens of replicate measurements. Under the higher light conditions, PinkyCaMP maintained strong baseline fluorescence (**Figure 3l**). Aside from significantly brighter fluorescence, PinkyCaMP demonstrated a higher ΔF/F at 10 Hz stimulation than RCaMP3 (**Figure 3n**), and a superior SNR of 63.5 ± 7.9 compared to RCaMP3’s 33.7 ± 4.8 (**Figure 3o**).

### Combined dual color fiber photometry for PinkyCaMP and sDarken

Next, we assessed the in vivo performance of PinkyCaMP. We chose to express PinkyCaMP in pyramidal neurons of the prefrontal cortex due to their well-established role in innate avoidance and decision-making behaviors^30^. We transduced the medial prefrontal cortex with either rAAV2/9.CamKII-PinkyCaMP or rAAV2/9.CamKII-mCherry and performed fiber photometry during an aversive air puff stimulus (**Figure S8**). PinkyCaMP exhibited robust Ca^2+^ transients in response to the air puff, clearly visible in individual traces. In contrast, control mice expressing only mCherry showed no detectable transients (**Figure S8d-g**). Significant fluorescence changes were observed in the PinkyCaMP group before and during the air puff (**Figure S8g**, p < 0.0001, ordinary one-way ANOVA), whereas no air puff-related changes were detected in the control group (p > 0.999, ordinary one-way ANOVA). Subsequently we determined whether PinkyCaMP could also indicatean approach-avoidance conflict in an elevated zero maze (EZM). As before, either PinkyCaMP or mCherry control was virally expressed in the mPFC (**Figure S9**). Consistent with previous reports, we observed that Ca^2+^ activity was lowest in the closed arms and increased when the animals transitioned to the open arms. PinkyCaMP-expressing mice showed a significant increase in fluorescence when transitioning from the closed to the open arm (**Figure S9**) (p < 0.01, ordinary one-way ANOVA), and a significant decrease when moving back to the closed arm (p < 0.01, ordinary one-way ANOVA).

To demonstrate the capability of PinkyCaMP in dual-color fiber photometry, we aimed to simultaneously image the innate avoidance response observed in our previous experiments, along with serotonin release in response to an aversive air puff. We co-expressed rAAV2/9.CamKII-PinkyCaMP together with a serotonin biosensor rAAV2/9.hSyn-sDarken^31^, or rAAV2/9.CamKII-mCherry with rAAV2/9.hSyn-nullmutant-sDarken, in the prelimbic cortex (**Figure 4a,b**). As expected, aversive air puffs elicited robust Ca^2+^ transients in pyramidal neurons, visible even in individual traces (**Figure 4c**). We observed consistent responses to the air puff (**Figure 4d,e**), with a significant increase in PinkyCaMP fluorescence during the stimulus (p < 0.001, ordinary one-way ANOVA), while mCherry showed no fluorescence changes (p = 0.9973, ordinary one-way ANOVA) (**Figure 4f**). Additionally, we detected serotonin release, as indicated by a decrease in sDarken fluorescence, coinciding with the increase in neuronal activity (**Figure 4g,h**). sDarken fluorescence was significantly reduced during and shortly after the air puff (p = 0.0001, ordinary one-way ANOVA), whereas the null mutant of sDarken exhibited no significant fluorescence changes before or during the air puff (p = 0.9119, ordinary one-way ANOVA) (**Figure 4j**).

**Figure 4:**
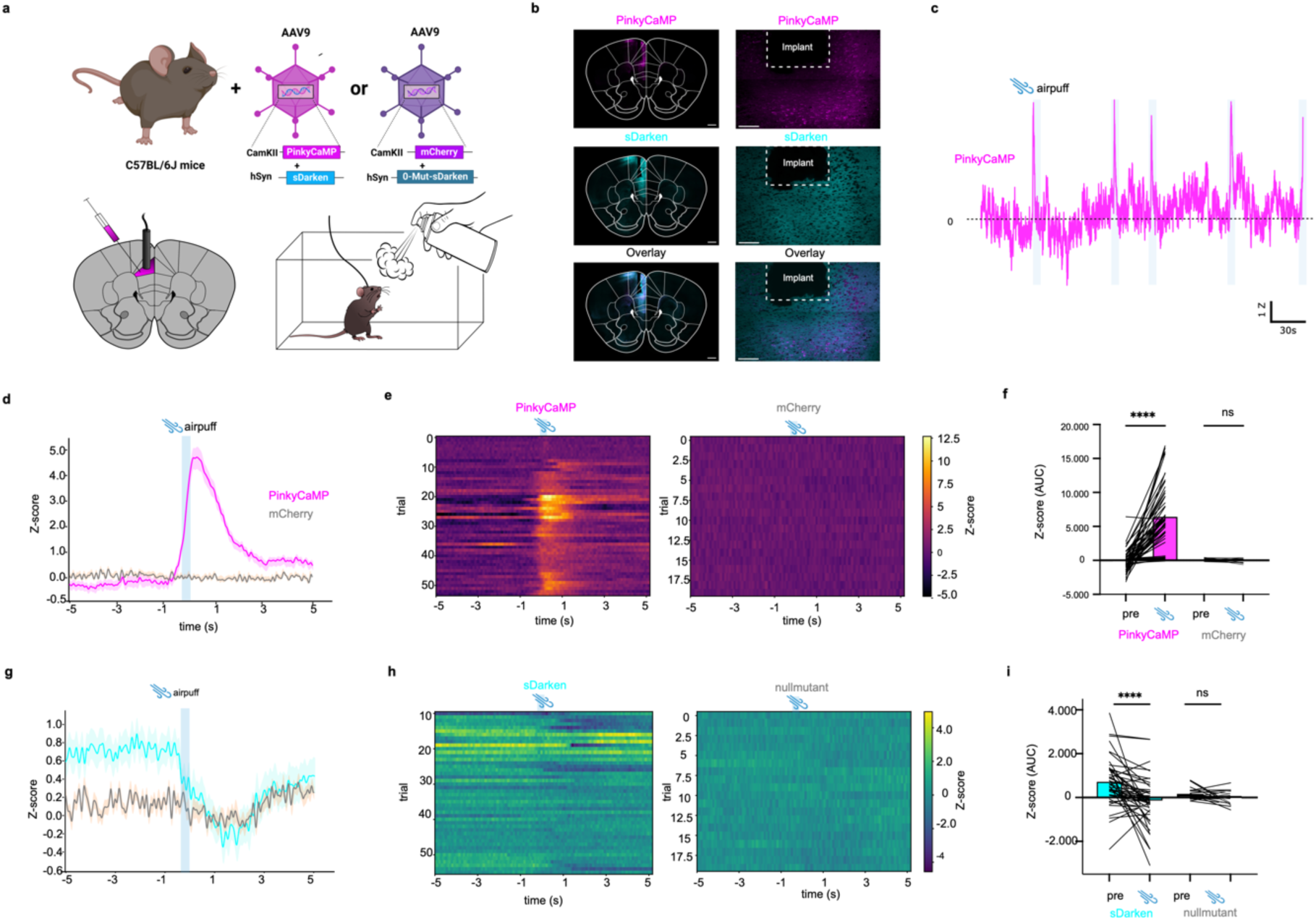
In vivo fiber photometry: Simultaneous imaging of neuronal activity with PinkyCaMP and serotonin with sDarken. (a) Schematic drawing of AAV injection into the prelimbic area (PrL) of the prefrontal cortex. Lower panel experimental setup for airpuff. (b) Histology example of PinkyCaMP expression and fiber placement. Scalebar 500 µm, magnification inset scalebar 100 µm. (c) Example traces of PinkyCaMP fluorescence in freely moving mice during an aversive airpuff in their homecage (airpuff). (c) Averaged PinkyCaMP activity aligned to an aversive airpuff n=7 mice (PinkyCaMP), n=4 mice (control)(mean ± s.e.m.). (d) Single trial heatmap of PinkyCaMP (54 trials from 7mice) and control (2 0 trials from 4 mice) (e) Area under the curve (AUC) of PinkyCaMP and mCherry signal before and during the airpuff. Ordinary one-way ANOVA, PinkyCaMP: mean pre 136 ± 211; mean during: 6503 ± 579; mCherry: mean pre 124 ± 44; mean during: -47 ± 64, **** p≤0.0001. (g) Averaged sDarken activity aligned to an aversive airpuff n=7 mice (sDarken), n=4 mice (nullmutant l)(mean ± s.e.m.). (h) Single trial heatmap of sDarken (54 trials from 7mice) and nullmutant (20 trials from 4 mice) (i) Area under the curve (AUC) of sDarken and nullmutant signal before and during the airpuff. Ordinary one-way ANOVA, sDarken: mean pre 749 ± 163; mean during: -170 ±165; nullmutant: mean pre 169 ± 59; mean during: -27 ± 67, **** p≤0.0001.

### PinkyCaMP is suitable for *in vivo* experiments in combination with blue light sensitive opsins

One caveat of red-emitting GECIs is that the 488 nm light-driven positive photoswitching can be interpreted as an increase in the GECI’s signal^32^. That is especially concerning in experiments in which the 488 nm light is directed to the same location where the red-emitting GECI is expressed. In order to test this, we designed an experiment to drive neuronal activity in principal cells via optogenetic disinhibition. Given the strong inhibitory input that granule cells (GC) receive from the dentate gyrus (DG) receive^33^, as well as the large proportion of GC that is silent^34–36^, we expected a strong increase in GC activity upon silencing of GABAergic interneurons. To this end, we injected the left DG of vGAT mice with rAAV2/9.CamKII-PinkyCaMP together with either the soma-targeted chloride-conducting opsin GtACR2 (rAAV/DJ.dlox-GtACR2-ST-mCerulean)^37^ or rAAV2/9.DIO-EGFP as control (**Figure 5a)**.

**Figure 5:**
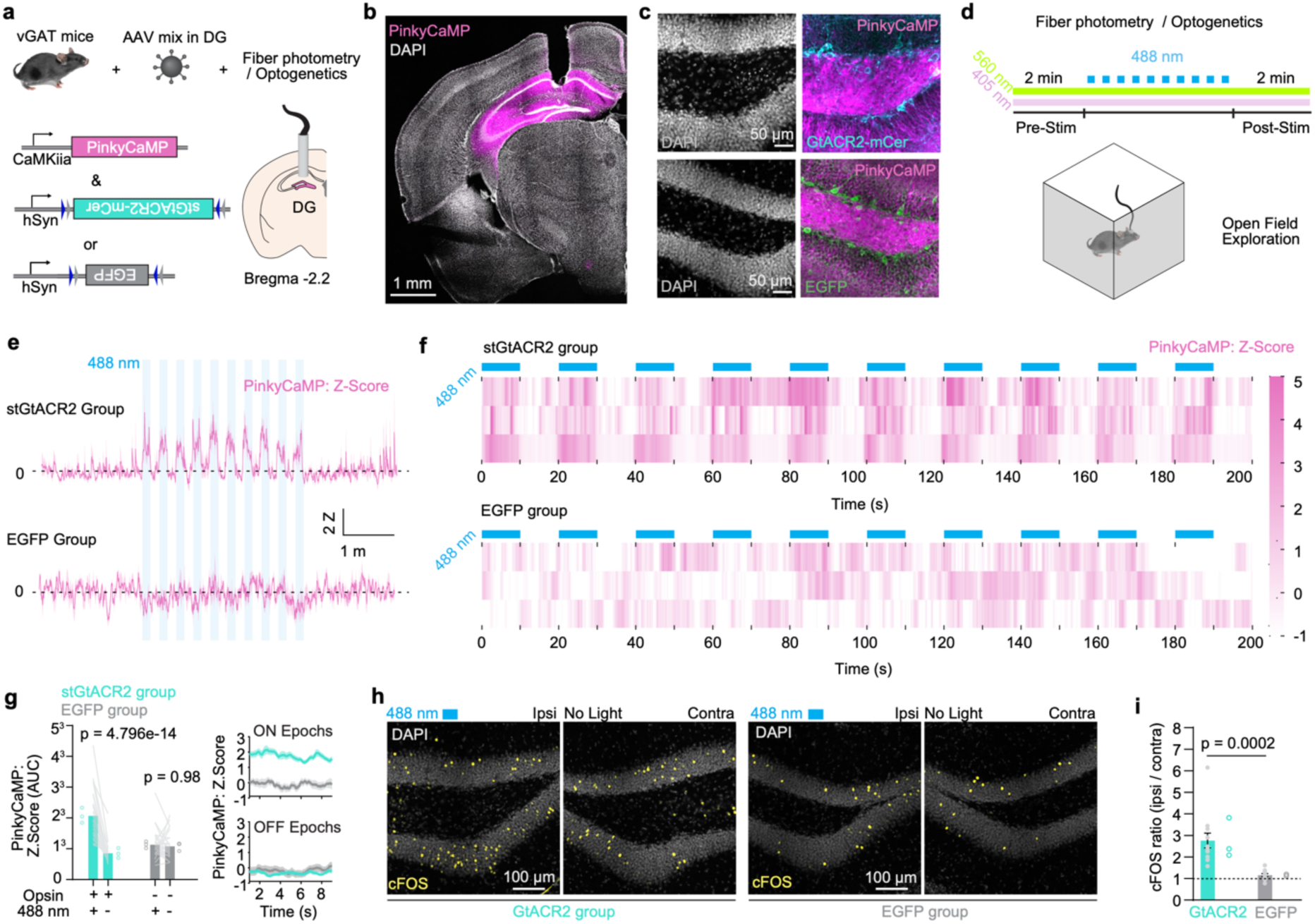
Combining PinkyCaMP with blue-sensitive optogenetics. (a) Experimental design. Fiber photometry recording of granule cell (GC) activity with PinkyCaMP. To drive GC activity, stGtACR2-mCerulean was expressed on vGAT+ neurons. EGFP was used as control. Exemplary immunofluorescent confocal images of (b). Fiber location and PinkyCaMP expression targeting dentate gyrus (DG). (c) Magnification of the hilar and GC layer regions of DG. DAPI (left), AAV-transduced neurons (right). vGAT expressing stGtACR2 (top) and EGFP (bottom). (d) PinkyCaMP fiber photometry during open field (OF) exploration. PinkyCaMP transients were measured using a 560 nm LED excitation light and a quasi isoemissive wavelength (405 nm) was used to control for motion artifacts. To test for positive photoswitching of PinkyCaMP while simultaneously driving GC activity, 488 nm light was switched ON/OFF while mice explored the OF (10X, 10 s ON / 10 s OFF). (e) Average PinkyCaMP traces (Z-score) during OF exploration in stGtACR2 group (n = 3 mice, top) vs. EGFP group (n= 3 mice, bottom). Blue shading indicates the periods when 488 nm light was on. (f) Heatmaps of PinkyCaMP signals (Z-score) during 488 nm light optogenetic-driven disinhibition. Blue squares represent the 488 light ON epochs. Each row represents an individual mouse. stGtACR2 group (top), EGFP group (bottom). g. Area under the curve (AUC) of PinkyCaMP Z-score during 488 nm light ON vs. OFF epochs. 488 nm light significantly increases PinkyCaMP fluorescence do to GC disinhibition (stGtACR2 group) and not due to positive photoswitching (EGFP group) (Šídák multiple comparison test, Z Score (AUC) during ON vs. OFF epochs stGtACR2 group: p = 7.23×10-15; EGFP group, p = 0.9993). 8h) Exemplary immunofluorescent confocal images comparing cFOS expression in GCs after OF exploration and optogenetic GC disinhibition between ipsilateral and contralateral to 488 nm light irradiation. i. Quantification of cFOS+ GC and shown as a ratio between the 488 nm irradiated (ipsilateral) vs contralateral side. 488 nm light increased cFOS expression in the stGtACR2 group while no increase was seen in the EGFP group (Unpaired, two-tailed t test, p = 0.0002).

This viral mix approach successfully and orthogonally transduced GCs with PinkyCaMP and vGAT cells with the opsin or the control EGFP (**Figure 5b-c**). We recorded GC PinkyCaMP transients during open field (OF) exploration with fiber photometry using 560 nm light and 405 nm light as the isoemissive wavelength control. Additionally, a 488 nm laser was switched ON/OFF during the exploration (10 s ON, 10 s OFF, 10X) through the same fiber (**Figure 5c**). As expected, PinkyCaMP signals increased during the light-ON epochs in the stGtACR2 group and had no effect on the EGFP group (**Two-Way-ANOVA, 488 nm light X Opsin interaction, p = 0.0015) (**Figure 5e-f**). While in the stGtACR2 group PinkyCaMP signals during the 488 nm ON epochs were significantly higher than during the OFF epochs, that was not the case for the EGFP group (Šídák multiple comparison test, Z Score (AUC) during ON vs. OFF epochs stGtACR2 group: p = 7.23×10^-15^; EGFP group, p = 0.9993) (**Figure 5g**). Importantly, the lack of increase in the red fluorescent signal during the ON epochs in the EGFP group confirms that indeed 488 nm light does not lead to positive photoswitching of PinkyCaMP. Finally, we observed a 3-fold increase in cFOS expression in the ipsilateral DG in the stGtACR2 group while no difference in the EGFP group was observed (*** Unpaired, two-tailed t test, ratio cFOS (ipsi/contra), p = 0.0002) (**Figure 5h-i**). The cFOS increase suggests that the observed PinkyCaMP signals report optogenetically driven disinhibition of GC activity. Collectively, these results showcase how PinkyCaMP can be used in single fiber photometry experiments combined with blue-light sensitive opsins without photoswitching artifacts.

### *In vivo* two-photon and one-photon miniature microscope PinkyCaMP imaging

First applications of GCaMP-imaging have been instrumental to understand basic hippocampal functions, such as place cells and their stability^38^. Moreover, in combination with red fluorescent indicators expressed in subsets of inhibitory interneurons allowed to discriminate the activity of different neuronal subsets^39^. Red-shifted Ca^2+^-indicators might be useful to simultaneously carry out Ca^2+^-imaging in different neuronal populations or cell types. Here, we tested the properties of PinkyCaMP in hippocampal CA1 neurons in awake head-fixed mice. We performed stereotactic injections of rAAV2/9.CaMKII-PinkyCaMP into dorsal CA1 of the hippocampus. Hippocampal windows were implanted a week later and imaging started 6-weeks post injection. We recorded Ca^2+^ transients in awake head-fixed mice at 10 Hz sampling rate at 1040 nm excitation wavelength **(Fig. 6a,b)**. The field of view contained more than 300 neurons and Ca^2+^ transients were analyzed off-line^40^ **(Fig. 6b)**. Ca^2+^ transients were reliably detected in PinkyCaMP-expressing CA1 neurons **(Fig. 6c,d, Supplementary Movie 1)**. We analyzed parameters of Ca^2+^ transients and measured a mean Ca^2+^-transient amplitude of 41 ± 0.6% **(Fig. 6e)**. CA1 neurons exhibited a transient rate of 3.2 ± 0.2 per minute **(Fig. 6f)**. We selected 129 representative unitary PinkyCaMP transients and measured an average rise time of 260 ± 10 ms **(Fig. 6g,h)**. These data show reliable measurement of Ca^2+^ transients with PinkyCaMP in CA1 neurons of awake head-fixed mice.

**Figure 6:**
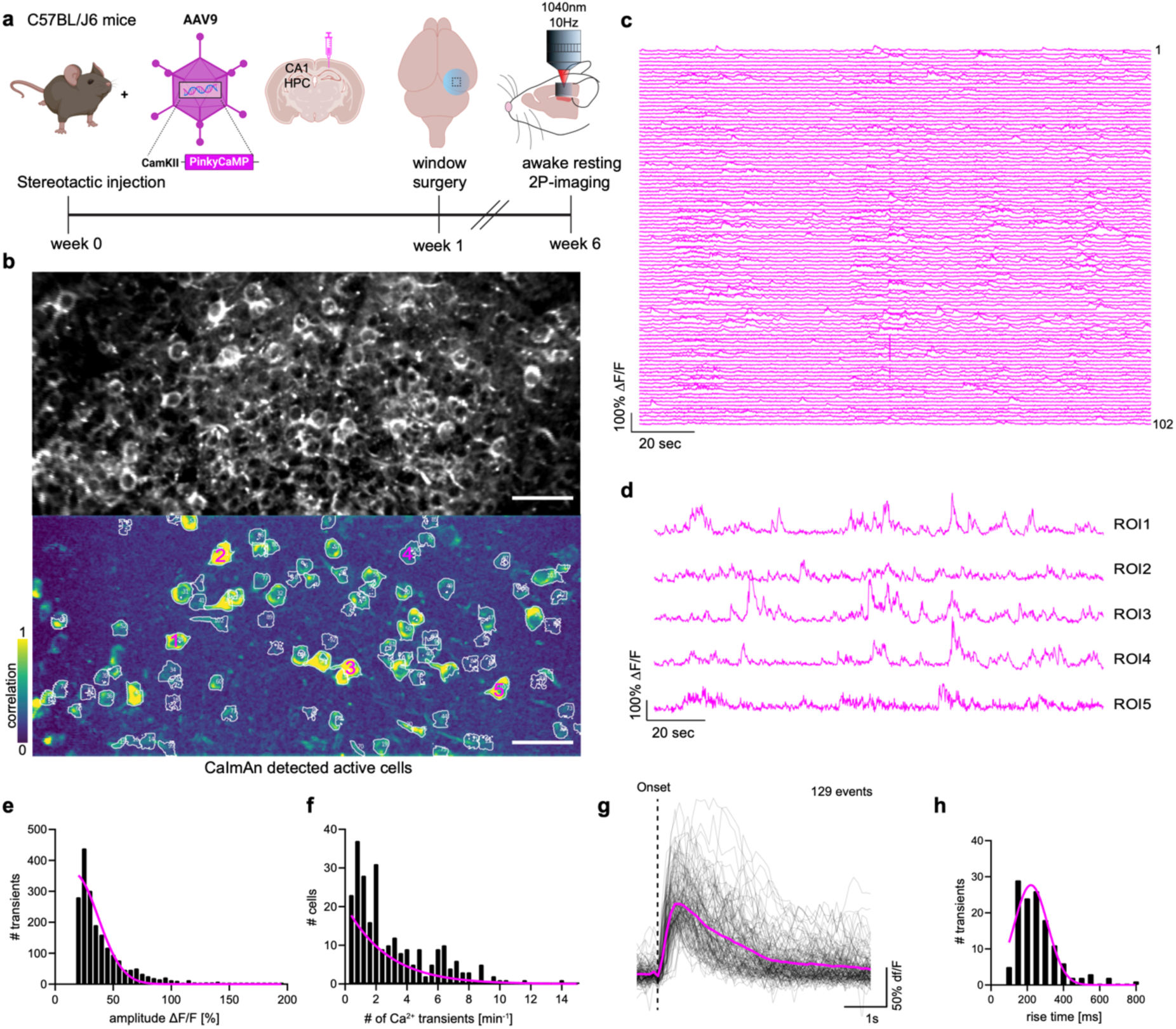
In vivo two-photon PinkyCaMP imaging. (a) Schematic of the experimental timeline for in vivo two-photon imaging. Stereotactic injection of rAAV2/9-CaMKII-PinkyCaMP into dorsal CA1 of the hippocampus at week 0, followed by window surgery during week 1. After 6 weeks of viral expression, awake head-fixed resting Ca2+-imaging was performed with a sampling rate of 10Hz at an excitation wavelength of 1040 nm. (b) Upper panel: Example of an imaging field of view (FOV) in hippocampal CA1, average intensity projected, scale bar 50 µm. Lower panel: Same FOV showing CaImAn identified cell footprints and the respective space correlation image. (c) Ca2+ transients of the cells detected in b (n=102 cells from one mouse). (d) Selected magnified Ca2+-traces of 5 cells from c. (e) Histogram showing the distribution of Ca2+ transients and their respective amplitude in % ΔF/F (mean = 41 ± 0.6%; n=2009 of 252 cells in 3 mice). (f) Number of cells over Ca2+-transient frequency (mean = 3.2 ± 0.2 min-1). (g) Average transient of 129 manually selected unitary Ca2+-events ordered by onset time (AvgAmp = 103 ± 3% ΔF/F). (h) Rise time histogram of cells in g (AvgRisetime = 260 ± 10 ms). Data are presented as mean ± SEM.

To assess PinkyCaMP’s compatibility with miniscope imaging, we tested it using a dual-color miniature microscope ( **Figure S10**). The experiment was performed analogous to the head-fixed two-photon imaging experiment (**Fig. 6**) with a gradient-index (GRIN) lens and the miniscope positioned above the hippocampal window. Ca²⁺ transients could be readily detected (**Supplementary Movie 2**).

## Discussion

In conclusion, we developed PinkyCaMP, the first mScarlet-based Ca²⁺ sensor. It bridges the performance gap between red-shifted Ca²⁺ sensors and green GECIs like GCaMP. PinkyCaMP is the brightest and most photostable red GECI, free from photoswitching, making it ideal for multicolor imaging and optogenetic compatibility. Compared to established red GECIs like jRGECO1a and jRCaMP1a^11^, it is an order of magnitude brighter with a superior signal-to-noise ratio, enabling effective low-light imaging and consistent long-term signal detection. Its photostability ensures reliable signal tracking, and its lack of photoswitching makes it an excellent choice for multiplexing, such as dual-color imaging or optogenetic experiments. However, we found that orange light excitation of PinkyCaMP can still cause residual activation of sensitive blue-light opsins, such as CoChR^26,27^. This could be circumvented by using more red-shifted excitation (> 590 nm) for PinkyCaMP or further blue-shifted channelrhodopsins could benefit multiplexing conditions^41–43^. As a red GECI, PinkyCaMP operates at wavelengths that typically result in lower tissue scattering, phototoxicity and autofluorescence, which will be in general advantageous for deep-tissue imaging^25,44,45^. In contrast to other R-GECO-based GECIs^7,11^, PinkyCaMP exhibited minimal lysosomal accumulation, highlighting its suitability for long-term expression without compromising neuronal health.

Notably, our experiments demonstrate that despite PinkyCaMP’s relatively slow kinetics, imaging pyramidal neuron and granule cell activity in the DG across various in vivo settings— such as fiber photometry, miniscope, and two-photon microscopy—is feasible. PinkyCaMP kinetics are comparable to jRCaMP1a, but it outperforms jRCaMP1a in terms of brightness, photostability and signal-to-noise. In the future, PinkyCaMP would benefit from modifications to support applications requiring faster temporal dynamics. The slow kinetics of PinkyCaMP that we observed is likely due to its high Ca^2+^ affinity. However, the current PinkyCaMP version seems ideal for neurons with sparse activity, where it will allow the detection of single events with high sensitivity. Additionally, PinkyCaMP has a relatively high sensitivity to calcium compared to other GECIs, which might lead to calcium buffering when overexpressed. This can likely be mitigated by carefully titrating the amount of virus and expression profile.

The next challenge will be to improve PinkyCaMP’s kinetics to match other GECIs. A feasible approach for this could be to exchange the RS20 peptide with either the ENSOP or ckkap peptide, as it has already been done for jGCaMP8^10^. Despite these limitations, PinkyCaMP enables easy imaging alongside other green GECIs or fluorescent biosensors in multi-color experiments and facilitates multiplexing with optogenetic tools. This advancement broadens the scope for analyzing interactions within neuronal populations, such as pyramidal cells and interneurons, as well as in various fundamental biological applications.

## Methods

### Animals

All experiments involving animals were carried out according to the guidelines stated in directive 2010/63/EU of the European Parliament on the protection of animals used for scientific purposes. All procedures involving animals were conducted in accordance with the guidelines of the responsible authorities and adhered to the 3R Principles.Experiments at the University of Bremen were approved by Senator für Gesundheit, Frauen und Verbraucherschutz of the Freie Hansestadt Bremen. Experiments carried out at the Ruhr University Bochum and at the DZNE, Bonn were approved by the Landesamt für Natur, Umwelt und Verbraucherschutz (LANUV). Experiments carried out at the Charité - Universitätzmedizin Berlin were approved by local authorities and the animal welfare committee of the Charité. Experiments performed at the University of Zurich were performed in compliance with the guidelines of the European Community Council Directive and the Animal Welfare Ordinance (TSchV 455.1) of the Swiss Federal Food Safety and Veterinary Office, and were approved by the Zürich Cantonal Veterinary Office.

For spectral multiplexing experiments with CoChR hippocampal neuronal cultures were prepared from P0 to P1 mice (C57BL/6NHsd; Envigo, 044) of either sex. For comparison with other red GECIs cortico-hippocampal slices were prepared from 7-9 day old CB57BL/6n mice of both sexes. Mice for post-mortem tissue removal were obtained from the Animal Facility of the Faculty of Biology and Biotechnology at Ruhr University Bochum, where they were bred and housed under standard conditions with food and water ad libitum under a 12 h light/dark cycle. For the dual color fiber photometry experiments and acute slice preparations male and female wild-type C57BL/6J mice (The Jackson Laboratory) were used. Animals were raised under standard 12-hour light/dark cycles and housed in groups in individually ventilated cages (IVC, Zoonlab) under controlled conditions (22°C ± 2°C, 50% ± 5% humidity). Food and water were available ad libitum. After surgical implantation, mice were individually housed. Experiments took place during the dark phase, aligning with the animals’ primary activity period. For optogenetic disinhibition of granule cells and PinkyCaMP fiber photometry recording, surgeries were performed on adult isoflurane-anesthetized vGAT mice (B6J.129S6(FVB)-Slc32a1tm2(cre)Lowl/MwarJ). For chronic in vivo two photon Ca2+ imaging and miniscope microscope imaging C57BL/6J mice were used.

### Molecular biology, protein purification, DNA constructs and availability of reagents

Phusion high-fidelity DNA polymerase (Thermo Fisher Scientific) was used for routine polymerase chain reaction (PCR) amplification. Restriction endonucleases, rapid DNA ligation kits, and GeneJET miniprep kits were purchased from Thermo Fisher Scientific. PCR products and products of restriction digests were purified using agarose gel electrophoresis and the GeneJET gel extraction kit (Thermo Fisher Scientific). DNA sequences were analyzed by DNA sequence service of Fasmac Co., Ltd.

To identify the initial prototypes, a portion of a tandem copy of the mScarlet gene, fused by a linker encoding GGTGGS, was amplified by PCR using three forward and three reverse primers. These primers were encoded for linkers of different lengths and had one NNK codon (**Figure S1a**). The product was digested by restriction enzymes BgIII and EcoRI. Starting from a plasmid encoding R-GECO1 in the pBAD vector, an EcoRI site was introduced into the RS20 to FP linker and a BglII site was introduced into the FP to CaM linker. This plasmid was digested with EcoRI and BgIII and the large fragment was ligated with the PCR product to create a library of PinkyCaMP prototypes. The ligation product was used to transform E. coli strain DH10B (Thermo Fisher Scientific) and 400 brightly red fluorescent colonies were picked from approximately 10,000 colonies inspected. Colonies illuminated by yellow light were screened visually using red-tinted goggles. The bacteria was grown in 1 mL of LB supplemented with 100 μg/mL ampicillin and 0.02% L-arabinose at 37°C overnight, then transferred to room temperature and incubated overnight. Proteins were extracted with B-PER (bacterial protein extraction reagent, Thermo Fisher Scientific) and the fluorescence brightness and the Ca^2+^-dependent response was assayed using a plate reader equipped with monochromators (Tecan). For the directed evolution of PinkyCaMP variants, libraries were generated by the overlap extension method(Ho et al. 1989). These libraries were screened as described above, and the most promising variants selected in each round were used as the templates for the next library creation.To construct the RCaMP3 plasmid, we first amplified the RS20 and cpmApple regions from a plasmid encoding R-GECO1.2, as well as the CaM region from jRGECO1a, with overlapping sequences. Subsequently, these two DNA fragments were combined by overlap-extension PCR. The full-length PCR product was then ligated into the pBAD vector using XhoI and HindIII restriction sites. Finally, the E217D mutation was introduced to generate RCaMP3. The genes encoding PinkyCaMP variants and RCaMP3 in the pBAD vector, which includes a 5’ poly-histidine tag, were expressed in E. coli. Cell pellets were lysed with a cell disruptor (Branson), and proteins were purified by Ni-NTA affinity chromatography (Agarose Bead Technologies).

The PinkyCAMP-CoChR bicistronic construct ( pAAV_CamKII(0.4)-PinkyCaMP-GSG-P2A-CoChR-ts-Kv2.1-HA-WPRE) was created by Gibson assembly^46^ using PinkyCamP, pAAV-hSyn-GCaMP6m-p2A-ChRmine-Kv2.1-WPRE (addgene #131004,ref^47^) and pAAV-EF1a-DIO-CoChR-Kv2.1-P2A-mScarlet-WPRE (Forli et al. 2021) as donors followed by restriction enzyme based sub-cloning (BamHI, HindIII) into a pAAV_CamKII(0.4) vector (addgene #198508, ref^48^). The Cre-dependent stGtACR-mCerulean AAV is available via the Viral Vector Facility of the University of Zürich. pGP-CMV-NES-jRCaMP1a (Addgene plasmid #61562), pGP-CMV-NES-jRGECO1a (Addgene plasmid #61563), pAAV-CaMKIIa-mCherry (Addgene plasmid #114469) and pAAV-Rep/Cap pAAV2/9n (Addgene plasmid #112865) are all available from addgene. All PinkyCaMP and sDarken constructs will be made available via Addgene (see https://www.addgene.org/Olivia_Masseck/) and via Viral Vector facility of the University of Zurich, or can be requested directly from the corresponding author.

### rAAV vector production

rAAV2/9.CamKII(0.4)-PinkyCaMP-GSG-P2A-CoChR-ts-Kv2.1-HA-WPRE was produced at the Charité Viral Core Facility (VCF, Berlin, Germany). All other adeno-associated viruses (AAV) were produced in house using the AAV helper-free vector system (Takara Bio). HEK cells (293tsA1609neo, Sigma Aldrich) were co-transfected with endotoxin-free plasmids: pHelper (Takara Bio), pAAV-Rep/Cap (pAAV2/9n (addgene plasmid #112865), and pAAV ITR expression vectors containing the gene of interest.

In 500 µL DMEM (Roti cell DMEM High Glucose, sterile, with glutamine, without pyruvate), 12 µg pHelper, 10 µg rep/cap plasmid, and 6 µg of the pAAV expression vector were mixed with 120 µL polyethylenimine (branched, MW ∼25000, Aldrich Chemistry), incubated at room temperature for 15 minutes, and dropwise added to dishes (TC-Dish 150 Standard, Sarstedt AG & Co. KG) containing HEK cells (70%-80% confluency). The media were replaced the following day with fresh media.

Three days post-transfection, supernatants were centrifuged (1500 g, 15 min), and AAVancedTM concentration reagent (System Bioscience) was added (4:1 ratio) to the cleared supernatants. After mixing, the solutions were incubated at 4°C for at least 12 hours. The mixtures were then centrifuged (4°C, 1500 g, 30 min), and the supernatants discarded. The pellets were resuspended in 500 µL DMEM + 10% FBS (Gibco), centrifuged (RT, 1500 g, 3 min), and the supernatants removed. The pellets were resuspended in cold, sterile 1x PBS.

For virus titer determination, 1 µL of the virus sample was incubated for 1 hour at 37°C with 120 µL DNase-digest buffer (10 mM Tris-Cl, 10 mM MgCl_2_, 50 U/mL DNase I [Roche Diagnostics]). The reaction was stopped by adding 5 µL EDTA solution (0.5 M) and incubating at 70°C for 10 minutes. The solution was then incubated at 50°C overnight with 120 µL proteinase solution (1 M NaCl, 1% N-lauroylsarcosine, 100 µg/mL proteinase K [Roche Diagnostics]). The following day, the solution was incubated for 10 minutes at 90°C, then 754 µL H2O was added. The sample was diluted (1:200) and virus titers determined by PCR using a standard curve of diluted plasmid samples.

### Measurement of biophysical properties in purified protein

Following purification the buffer was exchanged to 30 mM MOPS (pH 7.2) with 100 mM KCl. To perform pH titrations, protein solutions were diluted into buffers that contained 30 mM trisodium citrate, 30 mM sodium borate, 30 mM MOPS, 100 mM KCl, 10 mM EGTA and either no Ca or 10 mM CaCl_2_, and that had been adjusted to pH values ranging from 3 to 12. The protein solutions were mixed with buffers containing 30 mM MOPS (pH 7.2), 100 mM KCl, and various concentrations of free Ca^2+^ concentration(Tsien and Pozzan 1989). Absorbance spectra were measured using the Shimadzu UV1800 spectrometer and molar extinction coefficients were determined as previously reported(Cranfill et al. 2016). Fluorescence quantum yields were measured using a Hamamatsu Photonics absolute quantum yield spectrometer (C9920-02G) using an excitation wavelength of 560 nm.

Excitation and emission spectra were measured with protein solutions consisting of TBS with 10% glycerol (pH 8) containing either 10mM EGTA or CaCl₂ in a plate reader (Infinite 200 Pro, Tecan).

### Characterization of PinkyCaMP in HEK cell culture

Human embryonic kidney (HEK) 293T cells (tsA201 cells, ATCC) were cultured and transfected following the protocol described in ref^31^. Prior to imaging, cells were washed with TBS supplemented with 2 mM CaCl₂. Imaging and time-lapse acquisition were performed using an upright LNscope microscope (Lugis & Neumann) equipped with a 40x water immersion objective (LUMPLFLN40xW, Olympus) and a CMOS camera (ORCA-spark, Hamamatsu). To assess in vitro brightness, single images were captured using 560 nm illumination at 16 mW/mm². To evaluate photoswitching behavior, transfected HEK293T cells were imaged under the same conditions as described above at a frequency of 5 Hz. Continuous 560 nm illumination was applied throughout the recording. After a 30-second baseline period, a 470 nm laser (1 mW/mm², 10 ms pulses) was applied every 10 seconds for a total of six pulses.Photostability was assessed in transfected HEK293T cells under continuous illumination with 560 nm light at 1 mW/mm^2^.

### Primary dissociated hippocampal neuronal culture and gene delivery

Hippocampi were dissected and cells were dissociated by papain digestion followed by manual trituration. Neurons were seeded on glial feeder cells at a density of 1.6 × 10^4^ cells/cm^2^ in 24-well plates and maintained in Neurobasal-A supplemented with 2% B27 and 0.2% penicillin/streptomycin (Invitrogen). Neurons were transduced with rAAV (2.58 × 10^13^ viral genomes (vg) per well) at DIV 1-3 and were recorded between DIVs 14 and 21. Experiments using rAAV2/9.CamKII(0.4)-PinkyCaMP-GSG-P2A-CoChR-ts-Kv2.1-HA-WPRE were performed on an Olympus BX51 upright microscope equipped with a LUMPlanFL/IR ×40/0.80 W objective. Samples where constantly perfused (0.5 ml/min) with extracellular solution containing 140 mM NaCl, 2.4 mM KCl, 10 mM HEPES, 10 mM d-glucose, 2 mM CaCl2, 4 mM MgCl2 (pH adjusted to 7.3 with NaOH, 300 mOsm) 2µM NBQX, and 4µM Gabazine (SR-95531). Cell-attached measurements followed by whole-cell recordings where performed microelectrodes pulled from quartz glass capillaries (3–6 mΩ), filled with 136 mM KCl, 17.8 mM HEPES, 1 mM EGTA, 0.6 mM MgCl2, 4 mM MgATP, 0.3 mM Na2GTP, 12 mM Na2-phosphocreatine and 50 U ml−1 phosphocreatine kinase (300 mOsm), with pH adjusted to 7.3 with KOH. A Multiclamp 700B (Molecular Devices) and Digidata 1550B digitizer (both Molecular Devices) were used to control and acquire electrophysiological recordings as well as light engine LEDs, bipolar field stimulation and camera exposure timing. Electrophysiological data were acquired at 10 kHz and filtered at 3 kHz. Bipolar 1 ms, 66mA field stimuli were delivered through a Warner Instruments SIU-102 Stimulus Isolation to custom made electrodes (1 mm platinum wire) in the bath chamber.

For action spectra recordings, light from the Lumencor SpectraX23 light engine was filtered with narrow bandpass filters mounted on a FW212C filter wheel (Thorlabs) and delivered to the sample plane using a FM03R cold mirror (Thorlabs) in the epifluorescence beam path. The following filters were used (center wavelength ± 10 nm, Edmund Optics catalog no.): 372 nm (12147), 400 nm (65071), 422 nm (34496), 450 nm (65079), 480 nm (65084), 505 nm (34505), 535 nm (65095), 568 nm (65099), 600 nm (65102), 632 nm (65105) and 660 nm (86086). The light intensity for each wavelength was calibrated to the same photon flux corresponding to 0.325 mW mm^−2^ at 505 nm. Short light pulses were applied (1 ms) and the wavelength was either changed from UV to red light or vice versa or measurements were taken in both directions per cell. In the latter case, photocurrents were averaged per cell.

PinkyCaMP imaging was performed using a Semrock FF605-Di02 dichroic and a 620 LP ET longpass filter (Chroma) while the excitation light from the green-yellow LED of a Lumencor SpectraX 23 was filtered with a 586±15nm bandpass filter (FF02-586/15 Semrock). For stCoChR activation the blue LED of the SpectraX 23 filtered with a 438±15nm bandpass filter (438/29x Lumencor) was used.

Imaging was performed with a Hamamatsu ORCA-Fire digital CMOS camera (C16240-20UP) at 4×4 binning with 80 ms exposure at 10 Hz with 16 bit. Blue light application for stCoChR activation was performed with a duration of 1 ms between the imaging frames.

Light intensities were measured with a calibrated S170C power sensor (Thorlabs) at the sample plane. Electrophysiological data were recorded using Clampex 11.4, while imaging data were acquired using Micro-Manager (2.0)(Edelstein et al. 2014). Illumination spectra were recorded with an M-spectrometer (Thunder Optics, Montpellier, France). Electrophysiological data were acquired using Clampex.

### Organotypic slice cultures from mouse cortex and gene delivery

Slices cultures were prepared and cultured according to published protocols^49,50^. After separating the hemispheres, a parasagittal 45° cut was performed from the top of cerebral cortex to the center of the thalamus, the tissue placed in ice-cold Ringer’s solution (125 mM NaCl, 2.5 mM KCl, 1.25 mM NaH_2_PO_4_, 26 mM NaHCO_3_, 2 mM CaCl_2_, 1 mM MgCl_2_ and 20 mM D-glucose, saturated with 95% O_2_ and 5% CO_2_) and cut in 250-μm thick slices using a vibratome (Leica VT1200, 1 mm amplitude, 0.9 mm/s, 15° angle). The slices were temporarily stored in 34°C Ringer’s solution, until both hemispheres were cut. Under sterile conditions, the cortico-hippocampal slices were washed five times with HBSS (Sigma, H9394) and 2-to-3 slices were placed on one membrane insert (Millicell PICM0RG50, hydrophilized PTFE, pore size 0.4 μm). Slices were supplied with organotypic slice culture medium consisting of MEM (Sigma, M7278), 20% heat-inactivated horse serum (GIBCO/Life Technologies, 26050088), and additionally 1 mM L-glutamine, 0.001 mg/ml insulin, 14.5 mM NaCl, 2 mM MgSO_4_, 1.44 mM CaCl_2_, 0.00012 % ascorbic acid and 13 mM D-glucose and cultured at 37°C and 5% CO_2_. Transduction was performed at DIV 1 by adding 1 μl AAV suspension (up to 5-fold dilution in PBS (Sigma, 806552)) to the center of the cortex. A full medium exchange was performed every two days. Ca^2+^ Imaging experiments were performed at DIV13-22. Slices were placed in a custom recording chamber (1.5 ml) and superfused with Ringer’s solution at 1 ml/min at 24 °C using a peristaltic pump (Minipuls 3, Gilson) and an in-line heater, respectively. The chamber was placed under an upright microscope (Axioscope, Zeiss) fitted with a 10x/0.3 water immersion objective (W N-Achroplan, Zeiss). Epifluorescence excitation for all red fluorescent indicators was provided by a collimated 554 nm LED (MINTL5, Thorlabs) using a 560/40 nm excitation filter and a 585 nm dichroic mirror, while fluorescence was collected with a 630/75 nm emission filter (ET-TxRed filter set, Chroma). The light intensity in the focal plane was 0.1 mW/mm^2^ or 0.3 mW/mm^2^. GCaMP6f measurements were performed with a collimated 470 nm LED (M470L4, Thorlabs) using a 470/40 nm excitation filter, a 495 nm dichroic mirror (T495LPXR, AHF) and a 525/39 nm emission filter (BrightLine HC, Semrock). Here the intensity in the focal plane was 0.4 mW/mm^2^ or 0.7 mW/mm^2^. Images were acquired with a sCMOS camera (Orca-Flash 4.0 LT C11440, Hamamatsu) at 10 fps with 16-bit and 512 x 512 pixels (4×4 binning) using MicroManager 2.047. The slices were allowed to equilibrate for 10-20 min prior to imaging. Standard recordings lasted 10 min, long-term recordings lasted 60 min. Photoswitching recordings (**Fig. S6**) lasted 16 min with three 50-s intervals of additional high-intensity blue-light illumination in the focal plane (4.2 mW/mm^2^) followed by 200 s without blue light.

### In vitro electrophysiology in hippocampal slices

Hippocampal slices were prepared from 16 – 18-week-old Bl6 mice, three weeks after intracranial viral injection of either pAAV9-CaMKIIa-PinkyCaMP or control pAAV9-CaMKIIa-RCaMP3. Acute hippocampal slices were prepared as previously described (Whitebirch et al. 2022). In brief, sucrose-substituted artificial cerebrospinal fluid (dissection ACSF) and standard ACSF were prepared with filtered (0.22 µm) purified water. ACSF contained (mM): 22.5 glucose, 125 NaCl, 25 NaHCO_3_, 2.5 KCl, 1.25 NaH_2_PO_4_, 3 sodium pyruvate, 1 ascorbic acid, 2 CaCl_2_, and 1 MgCl_2_. Dissection solution used for slice preparation contained, in mM: 195 sucrose, 10 glucose, 25 NaHCO3, 2.5 KCl, 1.25 NaH_2_PO_4_, 2 sodium pyruvate, 0.5 CaCl_2_, 7 MgCl_2_. Dissection and standard ACSF were prepared freshly before each experiment and the osmolarity checked to range between 315 and 325 mOsm. Dissection ACSF was chilled on ice and bubbled with carbogen gas (95% O_2_ / 5% CO_2_, resulting in a pH of 7.3) for at least 30 mins before slice preparation. A recovery beaker was prepared with a 50:50 mixture of dissection and standard ACSF and warmed to 33°C. Mice were deeply anesthetized and transcardially perfused with ice-cold carbogenated dissection ACSF for approximately 30 – 45 sec. After decapitation, brains were quickly removed and transferred to ice-cold dissection ACSF in which the hippocampi were dissected free, placed into an agar block, and secured to a vibratome slicing platform with cyanoacrylate adhesive. Hippocampal slices were cut at 400 µm, parallel to the transverse plane. Slices were collected from the dorsal and intermediate hippocampus and transferred to the warm, continually carbogenated recovery beaker and allowed to recover for 30 min, after which the beaker was allowed to come to room temperature and left for an additional 90 min before start of the experiment.

Acute hippocampal brain slices were imaged in ACSF bubbled with carbogen gas (95% O₂, 5% CO₂) at 33°C and recorded at 50 Hz. Field stimulation was delivered using a glass patch pipette filled with 1 M NaCl, with its tip placed in the stratum oriens of CA2. Stimulation consisted of six 1 ms pulses of 300 µA, applied at varying frequencies using an isolated current stimulator (DS3, Digitimer Ltd.). PinkyCaMP fluorescence was recorded at low light intensity (0.23 mW/mm²) during 2 Hz and 5 Hz field stimulation.

### Surgeries

For stereotactic surgeries, viral injections and fiber-optic cannula implantation for dual color fiber photometry mice were initially anesthetized with 5.0% isoflurane in oxygen (1 L/min) and positioned in a stereotactic apparatus (Stoelting Co.) with integrated heating. Anesthesia was maintained with 1.5–2.0% isoflurane, monitoring respiration and toe pinch reflex. Carprofen (5 mg/kg) was administered subcutaneously for analgesia, and ophthalmic ointment (Bepanthen) applied to prevent corneal dehydration. After scalp shaving, the area was sterilized with ethanol and iodine, and lidocaine spray was applied. A midline incision was made, and the lambda-bregma distance measured. Holes were drilled above target regions. rAAV9-CaMKII-PinkyCaMP and rAAV9-hSyn-sDarken (or rAAV9-CaMKII-mCherry/AAV9-hSyn-0Mut-sDarken as control) were injected unilaterally into the right mPFC PrL at three depths (+1.70 mm AP, +0.30 mm ML, -1.85 mm, -1.75 mm, -1.65 mm DV). For ex vivo electrophysiology, rAAV9-CaMKII-PinkyCaMP, rAAV9-CaMKII-RCaMP3 were bilaterally injected into hippocampal CA1 (+2.10 mm AP, ±1.30 mm/±1.60 mm ML, -1.85 mm, -1.75 mm, -1.65 mm DV). Following injections, surgical sites were sutured (SMI, 191050).The skull surface was prepared with 37.5% phosphoric acid (Kerr, OptiBond™ FL kit) for up to 25 seconds to enhance implant adhesion. Ceramic fiber-optic cannulas (1.25 mm ferrule Ø, 400 µm fiber core Ø, NA: 0.5, 2.5 mm length) were implanted in the right mPFC PrL (+1.70 mm AP, +0.30 mm ML, -1.70 mm DV). The skull was coated with primer and adhesive (Kerr, OptiBondTM FL kit), UV-cured (850 W/cm²), and optical fibers were fixed with UV-cured dental cement (Geiz Dental GC 2278). After surgery, iodine ointment was applied to the skin, and mice recovered in clean, heated cages. Implanted animals were housed individually, and experiments began four weeks post-surgery to ensure viral expression.

For combining PinkyCaMP with blue sensitive optogenetics the left dorsal DG (-2.2AP, - 1.37ML, -1.9DV) was injected with a 1:1 virus mix of AAV9-CaMKii-PinkyCaMP-VariantC (9.8 x 10E12 vg/mL) and AAV-DJ.hSyn.chl.dlox.stGtACR2_mCerulean(rev).dlox-WPRE-hGHbp(A) (8.0 x 10E12 vg/mL) OR AAV9.hSyn.dlox.EGFP(rev).dlox-WPRE-hGHbp(A) (8.2 x 10E12 vg/mL). A single 400 nanoliter volume injection at a rate of 5 nanoliter per second was performed using a glass capillary and controlled by a microinjector (Nanoject III, Drummond Scientific). A self-made 400 µm core fiber with a 2.5mm metal ferrule cannula was implanted 0.1-0.2 mm above the injection site on the same surgery day. The ferrule was cemented (Tetric EvoFlow, A1) onto the skull. The mice had at least three weeks of recovery before the photometry recordings.

For chronic *in vivo* two photon Ca^2+^ imaging mice were anesthetized with an intraperitoneal (i.p.) injection of a mixture of Ketamine (0.13 mg/g) and Xylazine (0.01 mg/g) for a stereotactic injection of 1 μl of AAV9-CaMKII-PinkyCaMP into right dorsal hippocampus (+2 mm anterior-posterior, 2.3 mm lateral and -1.4 mm ventral, relative to Bregma) at 0.1 μl/min, using a UltraMicroPump, 34G cannula and Hamilton syringe (World Precision Instruments, Berlin, Germany).Stereotactic coordinates were taken from Franklin and Paxinos, 2008 (The Mouse Brain in Stereotaxic Coordinates, Third Edition, Academic Press). Analgesia was carried out with tramadol in the drinking water (0.1 mg/ml) for 3 consecutive days. Hippocampal window surgery followed one week after AAV injection and was performed as described before^51^. For head-fixation during in-vivo imaging a headpost (Luigs & Neumann, Ratingen, Germany) was cemented adjacent to the hippocampal window. Analgesia was carried out with tramadol in the drinking water (0.1 mg/ml) for 3 consecutive days. In vivo imaging was performed after 5 weeks of recovery. Meanwhile mice were habituated to rest head-fixed on a rotating disk in the microscope setup.

### Fiber photometry recordings for dual color imaging during an aversive airpuff and in the elevated zero maze

*In vivo* Ca^2+^ signals (PinkyCaMP) and serotonin dynamics (sDarken) were recorded using an RX10x LUX-I/O Processor and Synapse software (TDT). An integrated LED driver controlled three LEDs for excitation: 560 nm (Lx560, TDT) for PinkyCaMP and mCherry, 465 nm (Lx465, TDT) for sDarken and 0Mut-sDarken, and 405 nm (LX405, TDT) for the isosbestic control signal. Each wavelength was set to a light intensity of 25-30 µW and modulated at unique frequencies—530 Hz for 560 nm, 330 Hz for 465 nm, and 210 Hz for 405 nm. The LEDs were connected to a 6-port Fluorescence Mini Cube (Doric Lenses), with output delivered via a fiber-optic patch cord (NA: 0.48, 600 µm; Thorlabs) through a rotary joint (RJ1 1×1, Thorlabs) to a subject cable secured to the implanted ceramic fiber-optic cannula using an interconnect (ADAL2, Thorlabs). Emitted fluorescence was collected through the same fibers, separated by the filter cube’s dichroic mirrors, and detected by integrated photosensors at a sampling frequency of 1017.25 Hz. Event timestamps for airpuff applications were within the TDT system.

The airpuff, a robust anxiogenic stimulus, elicits a startle response in mice. Testing was performed in each mouse’s home cage under bright lighting conditions. Prior to testing, all cage enrichments (e.g., house, nesting material, paper roll) were temporarily removed and promptly returned afterward. Mice were allowed 5 minutes to habituate to the light, experimental room, and adjusted cage setup. A total of eight airpuffs were administered at intervals of at least 1 minute using compressed air directed toward the mouse. Airpuff timestamps were recorded and analyzed offline.

The elevated zero maze (EZM) had a white plastic floor with black, infrared-permeable walls surrounding the two closed areas. It featured a 55 cm inner diameter, 45 cm height, and 17 cm wall height. The test leverages mice’s natural exploratory drive in novel spaces and their aversion to open, elevated areas. The experiment was conducted under bright lighting with additional infrared LED illumination. Mice were initially placed at the entrance of a designated closed area (closed area A) and allowed to explore the maze for 15 minutes or until they made at least eight transitions between open and closed areas. If a mouse remained in closed area A for the entire session, it received a 5-minute break in its home cage, with the subject cable attached. For the second session, mice were placed at the entrance of the opposite closed area (closed area B) and allowed to explore for 15 minutes or until eight transitions were completed. This protocol effectively encouraged exploratory behavior. Transitions between open and closed areas were tracked offline.

### Photometry recording + optogenetics disinhibition of GCs

Fiber photometry recordings in the dorsal DG were performed using an iFMC6_IE(400-410)_E1(460-490)_F1(500-540)_E2(555-570)_F2(580-680)_S photometry system (Doric Lenses) controlled by the Doric Neuroscience Studio v6.1.2.0 software. A low-autofluorescence patch cord (400 μm, 0.57 N.A., Doric Lenses) was attached to the metallic ferrule on mouse’s head and used to excite PinkyCaMP with 560 nm (30 µW at the patchcord tip – 1 mW/mm^2^ irradiance) while collecting fluorescence emission measured by a photodiode detector (Newport). 405 nm was used as an isoemissive control fluorescence signal. Signals were sinusoidally modulated at 208 Hz and 333 Hz (405 nm and 560 nm, respectively) via lock-in amplification, then demodulated on-line and low-passed filtered at 12 Hz. Mice were connected to the patchcord 5 mins before the open field (OF) exploration in a new cage. Mice were placed in the center of the OF arena (50 x 50 cm) for 7.5 minutes while PinkyCaMP transients were recorded. 488 nm light (laser diode, 480 µW at the patchcord tip – 3.8mW/mm^2^) was alternated (10x 10 s ON / 10 s OFF) after 2 mins from the start of the PinkyCaMP photometry recording in the OF. After optogenetic silencing, mice explored the OF arena for 2 more mins. A tailored Matlab code was used to extract, process, and analyze PinkyCaMP signals.

### In vivo two photon Ca^2+^ imaging

Two-photon imaging was performed using an upright Zeiss 7 multiphoton microscope, equipped with an Insight x3 tunable laser (Spectra-Physics) and a 10x 0.5 NA objective (TL10X-2P, Thorlabs). The laser was operated at 1040 nm for PinkyCaMP fluorescence excitation, emission was isolated using a band-pass filter (617/73) and detected using a nondescanned detector. Zen blue software was used for microscope control and image acquisition. Image series (512×200 pixels, ∼460×189 μm field of view) were acquired at 10 Hz.

### Miniature microscope imaging

Miniature microscope images were acquired with a dual color miniature microscope (nVue 2.0, Inscopix) similar to the head-fixed two photon imaging. A GRIN lens (1 mm x 4 mm, Inscopix) was placed above the hippocampal cranial window and the miniature microscope was placed above the GRIN lens. Ca^2+^ traces were extracted with IDPS (Inscopix). Briefly, Ca^2+^ imaging videos were cropped around the imaging area and 4 times spatially down sampled (cutoff: 0.5, 0.005). Videos were motion corrected in IDPS^52^. Regions of interest (ROI) were selected manually and projected on the dF/F movie and calculated as the mean of the ROI / frame in IDPS.

### Histology

After ex vivo electrophysiology, acute brain slices were placed in 4% PFA at 4°C overnight, rinsed in 1x PBS, and mounted on Superfrost™ slides using Imaging Spacers (SecureSeal™, Grace Bio-Labs). Coverslips were applied with DAPI-containing mounting medium. All slides were stored at 4°C until fluorescence microscopy.

Following behavioral dual color fiber photometry experiments, animals received a lethal intraperitoneal injection of a ketamine/xylazine cocktail (130 mg/kg ketamine, 10 mg/kg xylazine) and were transcardially perfused with 1x phosphate-buffered saline (PBS), followed by 4% paraformaldehyde (PFA). Brains were extracted, stored overnight at 4°C in 4% PFA, and then transferred to a 30% sucrose solution before sectioning. Brains were frozen, and 45 µm coronal sections of target areas were cut using a cryostat. Sections were rinsed in 1x PBS, mounted on Superfrost™ slides (Thermo Scientific), and coverslipped with DAPI-containing mounting medium (ROTI Mount FluorCare with DAPI, Carl Roth GmbH).Viral injection sites and fiber-optic cannula placement were confirmed histologically using a confocal microscope (LSM880, Carl Zeiss) with Zen software. Images were acquired at either 10x or 20x magnification and processed in ImageJ.

Following optogenetic disinhibition of GCs 50 µm thick coronal brain sections were collected in 1X PBS using a vibratome (VT1200 S, Leica Biosystems). Four slices were selected in a range of 400 µm from the fiber location. Slices were blocked for 2h at room temperature (RT) in a solution containing 5% bovine serum albumin and 0.3% Triton in PBS). Transduced vGAT neurons were detected by incubating a primary antibody (AB) against GFP which recognizes both the EGFP and mCerulean (chicken a-GFP, 1:1000; catalog no. GFP-1010, Aves Labs). Additionally, a primary AB against cFOS (rat a-cFOS, 1:1000; catalog no. 226 017, Synaptic Systems) was used. Both primary ABs were incubated at 4°C overnight. The sections were washed (3x 10 min, 1x PBS) and incubated with secondary ABs (Alexa Fluor 488 donkey anti-chicken, 1:500; catalog no. 703-545-155, Jackson laboratories and Alexa Fluor 647 goat anti-rat, 1:500; catalog no. 31226, Invitrogen, conjugated in-house) for 3h at RT. Finally, the sections were washed again (2x 10 min, 1x PBS) and incubated with 4′,6-diamidino-2-phenylindole (DAPI, diluted in blocking solution with factor 1:10’000, 1x 10 min) before mounting on microscope slides using Hydromount (catalog no. HS-106, National Diagnostics).

A 2×2 tiled image of each dentate gyrus was taken from both hemispheres in all sections at a confocal laser scanning microscope (Axio Imager LSM 800, Zeiss) using a 25x oil-immersion objective (i LCI Plan-Neofluar 25x/0.8 IMM Korr DIC M27, Zeiss). Four z-stacks tile images at a resolution (pre-stitching) 1024×1024 pixel resolution were acquired. The Smart Setup function of the ZEN microscopy software (version 2.6, blue edition) was used to determine the optimal acquisition settings for the fluorophores used (DAPI, Alexa Fluor 488 (vGAT neurons), mScarlet (PinkyCaMP) and Alexa Fluor 647 (cFOS)). Each channel was acquired sequentially. For the overview image of PinkyCaMP expression, a 10x air objective (Plan-Apochromat 10x/0.45 M27, Zeiss) was used. Only DAPI and mScarlet channels were used and no z-stack was acquired at 1024×1024 pixels per image tile.

### Data analysis, quantification, statistic and reproducibility

To determine biophysical properties of the purified protein fluorescence intensity as a function of pH was then fitted by a sigmoidal function to determine the pKa. For Kd measurements. Fluorescence intensities versus [Ca2+] were plotted and fitted by a sigmoidal function to calculate the apparent Kd value for the purified protein.Calcium imaging data obtained in HEK cells were analyzed using ImageJ(Schindelin et al. 2012). Regions of interest (ROIs) were drawn around individual cells in the field of view, with an additional ROI used to measure the background fluorescence. The fluorescent brightness of each cell was calculated by subtracting the mean gray value of the background ROI from that of the cell ROI. Analysis was performed in ImageJ, with ROIs drawn as described above. ΔF/F values were calculated using the formula: ΔF/F= (F-F0)/F0 where F0 represents the fluorescence intensity of the first frame, and F represents the fluorescence intensity at a given time point. Statistical analysis was conducted in Python using a one-way ANOVA and Tukey’s post hoc test. To asses brightness fluorescence values were background-subtracted and normalized to the peak fluorescence for each respective sensor. The half-decay time (τ1/2) was determined by fitting a single-exponential decay curve to the mean fluorescence trace and extrapolating the time point at which the fluorescence decreased to half its initial value. All calculations were performed using Python. For multiplexing experiments ΔF/F Ca2+ traces were extracted from manually selected ROIs using the following equation: (F – F0)/F0 where F0 is the 3s median fluorescence before the first stimulus (field stimulation or photoexcitation of CoChR) is applied. Stimuli (type and pulse number) were performed randomly. Statistical analysis was performed with GraphPad Prism 10. 11 Electrophysiological data were analyzed using Clampfit 11 (both Molecular Devices). Cells were excluded from the analysis if the access resistance was above 25 MΩ or if the holding current exceeded 250 pA at -70mV holding potential. Cells were always patched randomly without any preselection by fluorescence. Calcium imaging data were analyzed using ImageJ(Schindelin et al. 2012). Briefly, ΔF/F Ca2+ traces were extracted from manually selected ROIs using the following equation: (F – F0)/F0 where F0 is the 3s median fluorescence before the first stimulus (field stimulation or photoexcitation of CoChR) is applied. Stimuli (type and pulse number) were performed randomly. Statistical analysis was performed with GraphPad Prism 10.

For the analysis of Ca^2+^ signals in cortical slice cultures data were processed in FIJI 2.16.0 (Schindelin et al. 2012). The images were further binned to 256×256 pixels and transformed to 32-bit. First, a region of interest (ROI) was defined by drawing a single polygon across the part of the slice which appeared focused, showed clear fluorescence above background and responses during synchronous network activity. A second region showing only membrane from the inserts was defined to obtain the background signal. The mean intensity value of the background region at t=0 was subtracted from all pixels in all images across the whole movie. Then a baseline fluorescence (F0) image was obtained for each stack by averaging 10 images at an early time point that showed fluorescence close to baseline (minimal intensity). Last, using the ImageJ image calculator, the stacks were converted to ΔF/F0 stacks. For further analysis, the mean ΔF/F0 signal of the ROI was transferred to Clampfit 11.2 (Molecular Devices). Data were collected in Excel Professional Plus 2019 (Microsoft) and statistically analyzed and plotted in OriginPro 2023 (OriginLab Corporation). Not all data sets were normally distributed and Kruskal-Wallis ANOVA (p < 0.05) was performed on all data followed by a Dunn’s multiple comparison. Figures were assembled in CorelDraw 2018 (Corel Corporation). For each sensor data were obtained from at least four independent transductions. Movies, which showed a focus shift, movement, unusual event heterogeneity or baseline drifts were not analyzed further. We also excluded single slices with unusually high or low baseline fluorescence intensity (F0) deviating >10-fold from the mean. All slices showed ΔF/F0 changes, which report on global synchronous network activity (cp. **Figure 3a,b**). The measured intensity changes reflect bursts of epileptiform network activity and they were identified as separate events, if the signal returned to around 50% compared to baseline. The event frequency was determined by counting the number of events in the first 300 s of each movie. To avoid undersampling because of slow calcium dynamics or sensor kinetics, only slices with event frequencies between 0.05 to 0.25 Hz were taken into account (cp. **Figure S5f**).

Quantified brightness values give the mean F0 signal (background-subtracted, see above) of the ROI. The relative signal change is reported as the peak ΔF/F0 value of the first 20 events that were identified in each slice. The absolute signal strength of these events was calculated by multiplying their peak ΔF/F0 value with the mean brightness (F0) of the corresponding slice. Time constants ***τ*** of the signal decays were obtained by fitting single exponentials to the decay region. For determining the photostability (bleaching) in each slice (**Figure 3g**), the average baseline intensity (offset from exponential fits) after the first four events at the beginning of the recording and after four events close to 10 min recording time was determined. Then the obtained baseline intensities from the beginning were subtracted from the 10 min values and multiplied by 100 (to give %). Photoswitching (**Figure 3h**) describes the difference in baseline fluorescence intensity after blue-light illumination (measured within 0.1 s after the LED was turned off) compared to before blue-light illumination (see **Figure S6**). Here, up to three measurements were averaged per slice, however, stimuli that were directly followed by a synchronous event were excluded.

For acute brain slices regions of interest (ROIs) were manually drawn around visible somas and a random section of neuropil. For PinkyCaMP, the neuropil signal was scaled by a factor of 0.7 (as described previously53)before being subtracted from the somatic signal. To account for photobleaching and artifacts, an exponential decay was fitted to each ROI and subtracted, followed by applying a rolling median with a five-frame window. The maximum ΔF/F of each cell was recorded and averaged to compare responses at different stimulation frequencies. All data was statistically analyzed with a Shapiro-Wilk test and Mann-Whiney U test in GraphPad Prism.

RCaMP3, recordings required a much higher light intensity (11.83 mW/mm²). Both sensors were measured under these conditions during 2 Hz and 5 Hz stimulation. Due to RCaMP3’s neuropil signal magnitude exceeding that of the somas, neuropil subtraction was omitted to prevent the appearance of negative responses (data not shown).

For 5 Hz stimulation, we compared the maximum ΔF/F averaged across cells, brightness: calculated as the mean fluorescence of cells in the first frame and signal-to-noise ratio (SNR): was defined as the maximum ΔF/F divided by the mean standard deviation of the one-second pre-stimulus baseline between PinkyCaMP and RCaMP3. Statistical comparisons of these metrics were performed using Shapiro-Wilk tests for normality and Mann-Whitney U tests in GraphPad Prism.

For cFOS Image Analysis Bit-plane Imaris (Oxford instruments, version 10.2.0) was used to process, analyze and quantify granule cell (GC) cFOS expression. First, the surface creation tool was used to mask the GC layer visualized by DAPI. The parameters for surface creation were a surface grain size of 2 µm and a threshold for absolute intensity of the signal > 29.1 (a.u.). This surface was used to mask the cFOS channel, so as to restrict the cFOS positive (+) cells counted to the detected GC layer. Finally, the spot detection algorithm was used on the mask to automatically detect the number of cFOS+ cells. The parameters used were an estimated cell diameter of 8 µm and a quality above 11.8, where quality is a measure of signal intensity. The volume of the GC layer, as well as the number of detected cFOS spots were extracted and further analyzed using R on R-studio (ver. 4.0.4 and 1.4.1106 respectively). To compute the number of cFOS+ GCs, the GC number per image was estimated using the reported numerical density^54^. The cFOS fraction was calculated by dividing the cFOS+ cells by the estimated GCs per image. Finally, cFOS was reported as a ratio between the 488 nm irradiated (ipsilateral) vs contralateral side.

For data analysis of dual color fiber photometry, a custom Python script was developed following the guidelines that have previously been outlined^55^. The analysis script is publicly available on GitHub (https://github.com/masseck/FibPho-PinkyCaMP.git).

For two-photon data analysis, recorded Image series were motion corrected using the python toolbox for Ca2+ data analysis CaImAn40 applying rigid-body registration. Detection of cell bodies and source-separation was performed using the CaImAn algorithm based on constrained non-negative matrix factorization56. ΔF/F Ca2+ traces were extracted from detected ROIs using the following equation: (F – F0)/F0, where F0 is the minimum 8th quantile of a rolling window of 200 frames. Ca2+ imaging traces were processed and analyzed using Gaussian process regression (GPR)57–59 to obtain a smooth approximation of the fluorescence signal over time. Peaks in the GPR-predicted traces were identified and characterized by fitting an exponentially modified Gaussian (EMG) function60. Metrics such as peak amplitude were extracted for each identified peak. Baseline fluorescence was calculated as the average of the 10th percentile of fluorescence values during the baseline window (10–45 s) For all following distribution analysis, a filter was applied to only select transients with >20% ΔF/F. For each cell the number of Ca2+ transients per minute (>20% ΔF/F) were calculated. To ensure reproducibility61, the analysis pipeline was implemented in Python and details of the computational environment and dependencies are provided. Additionally, 129 representative unitary events were selected by hand and extracted from ΔF/F Ca2+ traces with an amplitude of at least 60% ΔF/F using Igor Pro (Wavemetrics, Lake Oswego, OR, USA). Rise time was calculated from these events using the 10-90% time interval of onset until peak of the respective event transient.

Statistical analysis, and graph preparation were carried out using GraphPad Prism 9 (GraphPad Software Inc, La Jolla, CA, USA). To test for normal distribution of data, D’Agostino and Pearson omnibus normality test was used. Statistical significance for groups of two normally distributed data sets paired or unpaired two-tailed Student’s t-tests were applied. One-way ANOVA with Šídák’s multiple comparison test was performed on data sets larger than two, if normally distributed. If not indicated differently, data are represented as mean ± SEM. Chosen sample sizes were similar to commonly used ones in the community. We did not predetermine the sample size. Whenever possible automated data analysis was used.

## Acknowledgments

We would like to thank Celina Schreiber, Frederik Piel, Maja Neubauer, Monika Dopatka, Hannah Urbschat and the Charité Viral Core Facility for expert technical assistance and Jana Ottens for pre screening of cpmScarlet variants. O.A.M was funded by the Deutsche Forschungsgemeinschaft (DFG MA 4692/6-3, MA 4692/12-1). Work at The University of Tokyo was supported by grants from the Japan Society for the Promotion of Science to T.T. (KAKENHI 21H00273, 23H02101, and 24H00267) and to R.E.C. (24H00489, 24H02267, and 19H05633) and a grant from the National Institutes of Health (RF1NS126102 to PI Shy Shoham). S.I. was supported by the MERIT-WINGS program of the University of Tokyo. Work at Ruhr University Bochum was supported by the German Research Foundation (DFG; project 394431587 – FOR2795 to A.R.). Work at the Charité was supported by the German Research Foundation (DFG), project 184695641 – SFB 958, project 327654276 – SFB 1315, Clinical Research Unit KFO 5023 ‘BecauseY’ / Project number 504745852, project 415914819 - FOR 3004, project 431572356 and under Germany’s Excellence Strategy – Exc-2049-390688087), by the European Research Council (ERC) under the European Union’s Horizon 2020 research and innovation program (BrainPlay Grant agreement No. 810580), by the Federal Ministry of Education and Research (BMBF, SmartAge – project 01GQ1420B) and by the Einstein Foundation Berlin (grant ID EZ-2014-226). Work at the University of Zurich was supported by the European Research Council (ERC) under the European Union’s Horizon 2020 research and innovation program (grant agreement No. 891959 to T.P.) as well as by the Swiss National Science Foundation (project grant no. 310030_196455 to T.P.). N.G., F.F., M.M. and MF were supported by the European Union ERC-CoG (MicroSynCom 865618), the German research foundation DFG (SFB1089 C01, B06; SPP2395), and by the iBehave network to MF (funded by the Ministry of Culture and Science of the State of North Rhine-Westphalia). We would like to thank Jean Charles Paterna and the Viral Vector Facility of the University of Zürich for producing the Cre-dependent stGtACR-mCerulean AAV. pGP-CMV-NES-jRCaMP1a (Addgene plasmid #61562) and pGP-CMV-NES-jRGECO1a (Addgene plasmid #61563) were a gift from Douglas Kim & GENIE Project. Hanns Ulrich Zeilhofer provided the vGAT mouse line. pAAV-CaMKIIa-mCherry was a gift from Karl Deisseroth (Addgene plasmid #114469).

## Author Contributions

O.A.M. conceived and supervised the project. S.I. discovered the PinkyCaMP0.1a,b prototypes and performed all of the directed evolution to produce PinkyCaMP as represented in Figure 1 and Supplementary Figure 1, under the supervision of T.T. and R.E.C.. Together, R.F. (under the supervision of O.A.M.) and S.I. (under the supervision of T.T. and R.E.C.) characterized purified proteins. Together, R.F. and M.K. characterized all variants in HEK cells, brain slices and produced custom made viruses under the supervision of O.A.M.). J.W. designed the bicistronic PinkyCaMP CoChR with the help of A.K, performed, analyzed and interpreted all related Data (Fig. 2) under the supervision of D.S.. Organotypic slice experiments were performed by G.L., T.Z., R.F. and supervised by A.R. Brain slice recordings were done by R.F., V.K. and supervised by O.A.M. and S.H. Dual color fiber photometry experiments were performed and analyzed by K.R. and O.A.M. P.L.M Experimental design for in vivo photometry and optogenetics disinhibition, stereotaxic surgeries, photometry recording and analysis, manuscript writing & editing, under the supervision of T.P.. A.C. tissue preparation and immunofluorescence staining for photometry and optogenetic experiment, confocal imaging and image analysis under the supervision of T.P. In vivo 2-Photon measurements were performed and analyzed by N.G., F.F. and M. M. and supervised by M.F. All authors contributed equally to the data analysis and writing of the manuscript.

## Data availability

The authors declare that the data supporting the findings of this study are available within the paper, the methods section, and Extended Data files. Raw data will be deposited on open accessible repositories like DRYAD.

## Use of AI

During the preparation of this work, the authors used ChatGPT4o mini to improve english language and readability. After using this tool the authors reviewed and edited the content as needed and took full responsibility for the content of the publication.

