## Supplementary material for "PinkyCaMP a mScarlet-based calcium sensor with exceptional brightness, photostability, and multiplexing capabilities"

### Supplementary Figures

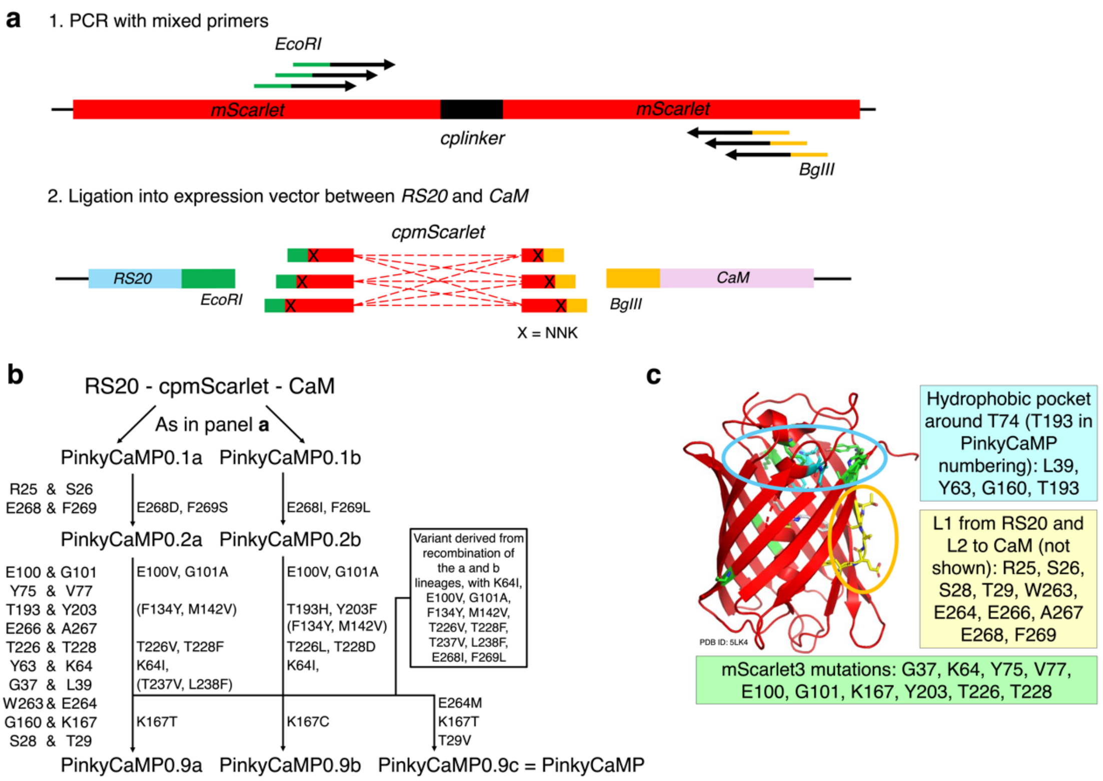


**Figure S1: The development of PinkyCaMP.** (**a**) Schematic representation of the method used to initially identify promising prototypes. We generated a library of circularly permuted mScarlet (cpmScarlet) variants with a calmodulin (CaM)-binding peptide (RS20, a variant of CaM-binding region of the smooth muscle form of myosin light chain kinase, derived from R-GECO1)[6] fused to the N-terminus and CaM (also derived from R-GECO1) fused to the C-terminus. This library was constructed such that the linker connecting cpmScarlet to RS20 was 2-4 residues in length, and the linker connecting cpmScarlet to CaM was 5-7 residues in length. The gene encoding cpmScarlet with various linkers was ligated into a pBAD expression plasmid containing the 5' *RS20* gene followed by an *EcoRI* site and the 3' CaM gene preceded by a BglII site. The codons immediately 3' to the EcoRI restriction and immediately 5' to the BglII site were randomized using an NNK codon (where N = A, G, C, T and K = G, T). The theoretical diversity of this library was 9,216 gene sequences or 3,600 protein sequences. This library was expressed in the context of E. coli colonies and 400 of the brightest red fluorescent colonies were picked out of ~10,000 colonies visually inspected. These 400 clones were cultured, the protein was extracted, and evaluated in terms of brightness and response to Ca2+. Two promising prototypes were identified and designated as PinkyCaMP0.1a (brighter; ΔF/F ~ 1.1; RS20 to FP linker = RSW and FP to CaM linker = EGEAEF) and 0.1b (more responsive; ΔF/F ~ 3.6; RS20 to FP linker = RSW and FP to CaM linker = EPEAEF). (b) Lineage of PinkyCaMP variants. To further improve the performance, PinkyCaMP0.1a and PinkyCaMP0.1b were subjected to directed evolution. As demonstrated for other single FP-based biosensors, the linker regions (i.e., RS20 to FP and FP to CaM) have a particularly large impact on biosensor function. In the prototype biosensors, the linker regions include two residues for which the codons correspond to the EcoRI and BglII restriction sites (encoding for RS and EF, respectively). We first optimized these residues by using site-saturation mutagenesis to fully randomize the positions (NNK codons) and then performing colony based screening and Ca2+-response assays (0 and 39 μM) on crude protein extracts. For the RS20 to FP linker, R25 and S26 of both PinkyCaMP0.1a and 0.1b were randomized using the overlap extension method. The two resulting libraries were screened in parallel as described above, with evaluation of extracted proteins for 200 bright clones out of ~2,000 colonies examined for each. For the FP to CaM linker, the E268 and F269 residues were optimized similarly. The best performing variants, PinkyCaMP0.2a and 0.2b, both retained the original RS sequence in the RS20 to FP linker, and had the EF of the FP to CaM linker changed to GS and IL, respectively. The brighter PinkyCaMP0.2a exhibited a Ca2+-dependent ΔF/F ~ 1.7 and the more responsive PinkyCaMP0.2b exhibited a ΔF/F ~ 7.7 in crude lysate. To further improve the performance, we next focussed our attention on optimization of the cpmScarlet domain and systematically screened libraries created by randomization of 20 additional positions in the protein, two residues at a time. To optimize these 20 positions, starting from both PinkyCaMP0.2a and PinkyCaMP0.2b in parallel, ten iterative libraries were created in which two residues at a time were randomized with an NNK codon and screened as described above. The top one or two winning clones from each library were used as the template for the subsequent library generation. When optimizing G37 and L39 residues, we identified a variant that exhibited a better balance of brightness and response and so we designated it as a third “c” lineage. At the completion of this optimization process we had three high-performance variants designated as PinkyCaMP0.9a, PinkyCaMP0.9b, and PinkyCaMP0.9c. (c) The position of the 24 residues subjected to site-saturation mutagenesis mapped on the crystal structure of mScarlet (PDB ID : 5LK4).21 Ten of these 24 positions are the residues mutated during the development of mScarlet3 (G37, L64, Y75, V77, E100, G101, K167, Y203, T226, T228; colored green).23 Further inspired by the development of mScarlet3, which emphasized the hydrophobic pocket around T74 of mScarlet (corresponding to T193 of PinkyCaMP) we also focussed on the four hydrophobic pocket residues L39, Y63, G160, and T193 (colored cyan). Finally, to further optimize residues located at or close to the linker regions, we targeted R25 (not shown), S26 (not shown), S28 and T29 in the RS20 to FP linker and W263, E264, E266 (not shown), and A267 (not shown), E268 (not shown), and F269 (not shown) in the FP to CaM linker (colored yellow).

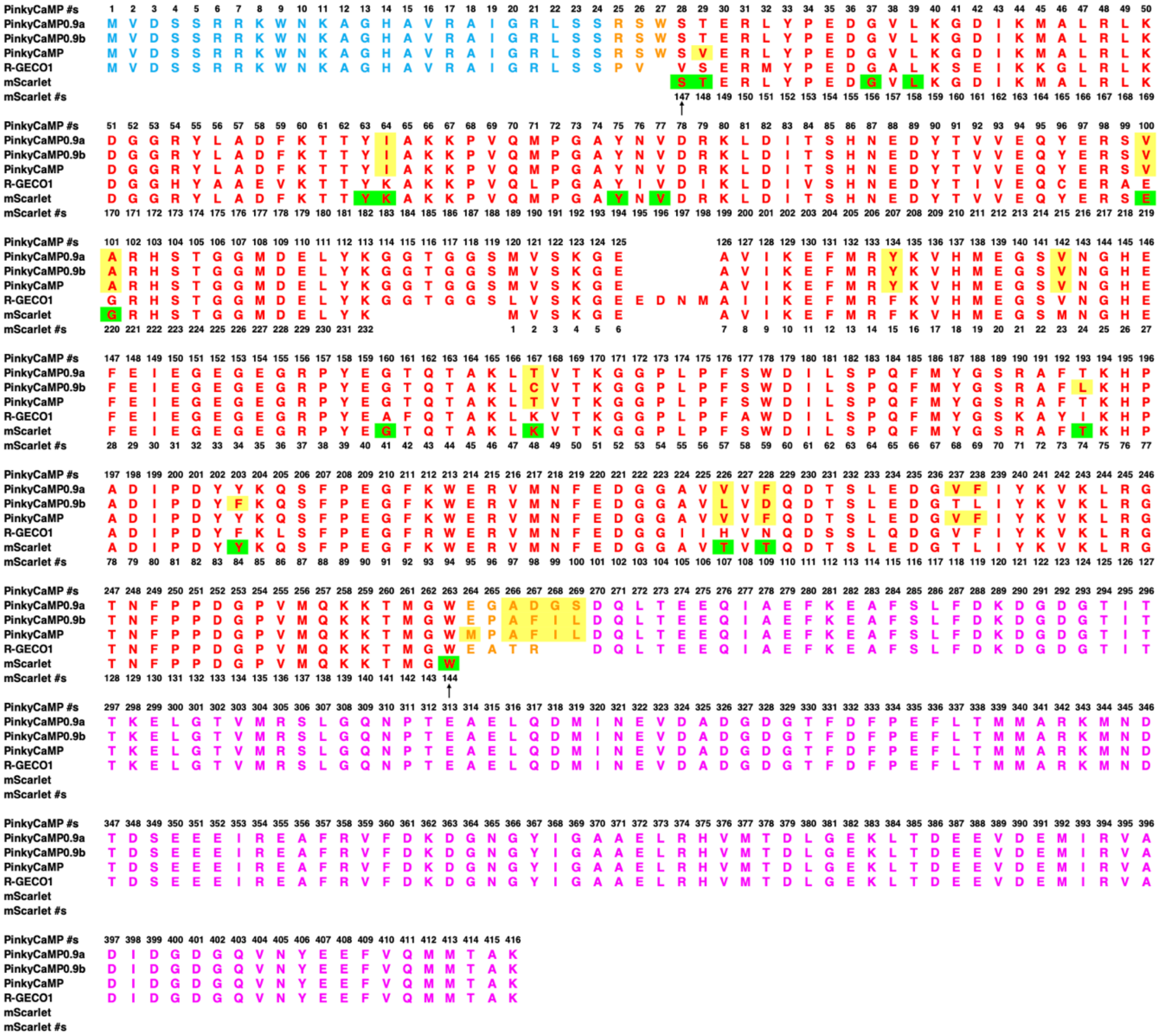

**Figure S2: Sequence alignment of mScarlet, R-GECO1 and PinkyCaMP variants.** The mScarlet gene sequence has been permuted in order to align with the cpmScarlet domain of PinkyCaMP and the mApple domain of R-GECO1. RS20 is colored in cyan, the FP domain is colored in red, CaM is colored in magenta and the linkers are colored in orange. The mutations accumulated during the optimization process are highlighted in yellow. The residues of mScarlet subjected to site-saturation mutagenesis are highlighted in green. During site-saturation mutagenesis, the original residue was identified as the preferred residue at some positions. The gate post residues are indicated by arrows.

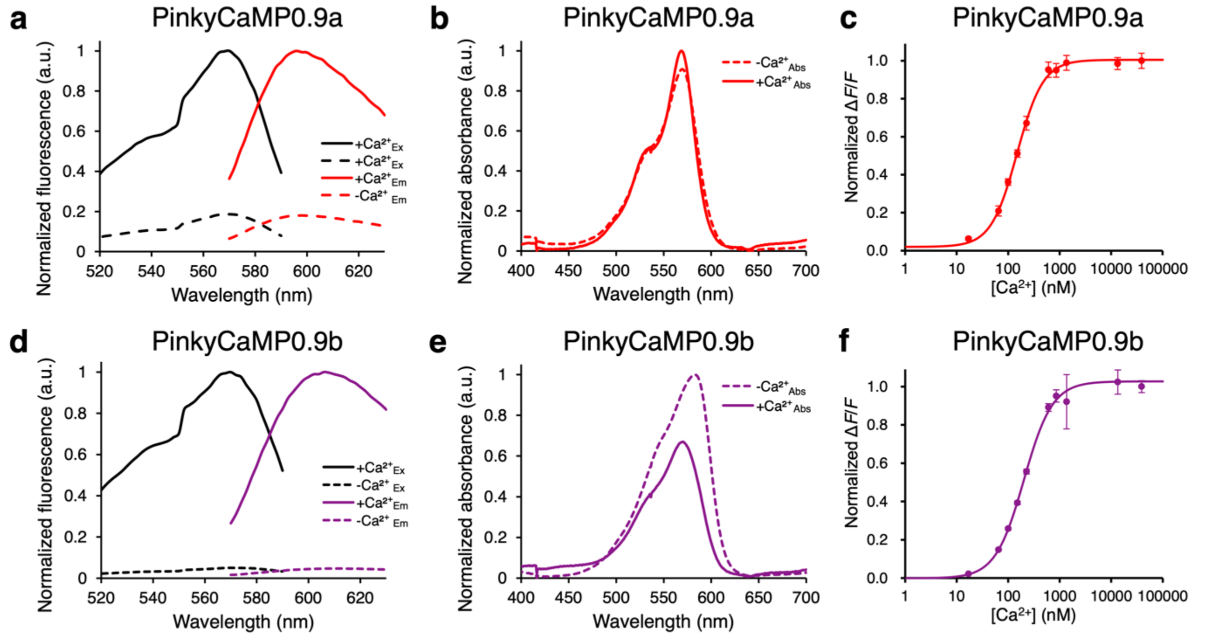


**Figure S3: *In vitro* characterization of PinkyCaMP0.9a (a-c) and PinkyCaMP0.9b (d-f).** The characterization of PinkyCaMP0.9c (= PinkyCaMP) is provided in Figure 1d,e,f. (a,d) Excitation (emission at 630 nm), and emission (excitation at 535 nm) spectra in the presence (39 μM) and absence of Ca2+ of PinkyCaMP0.9a (a) and PinkyCaMP0.9b (d). (b,e) Absorbance spectra in the presence (39 μM) and absence of Ca2+ of PinkyCaMP0.9a (b) and PinkyCaMP0.9b (e). (c,f) Ca2+ titration curve (n = 3 replicates; mean ± s.d.) of PinkyCaMP0.9a (c) and PinkyCaMP0.9b (f). To summarize, in the presence of Ca2+, PinkyCaMP0.9a has absorbance and emission peaks at 569 and 596 nm, respectively. It exhibits a Ca2+-dependent ΔF/F of 3.4 and has an apparent Kd for Ca2+ of 149 nM. The fluorescence response is primarily due to an increase in the quantum yield (0.16 to 0.44). The extinction coefficient remains fairly constant, only increasing from 89,000 M-1cm-1 to 98,000 M-1cm-1 upon binding to Ca2+. Relative to PinkyCaMP0.9a, PinkyCaMP0.9b exhibited slightly red-shifted absorbance and emission peaks at 570 and 606 nm, respectively. It has a ΔF/F of 21.9 and an apparent Kd of 202 nM. As with PinkyCaMP0.9a, the fluorescence response is entirely attributed to a large quantum yield change from 0.02 to 0.43, with the extinction coefficient actually decreasing from 85,000 M-1cm-1 to 57,000 M-1cm-1, upon binding to Ca2+ (Table 1).

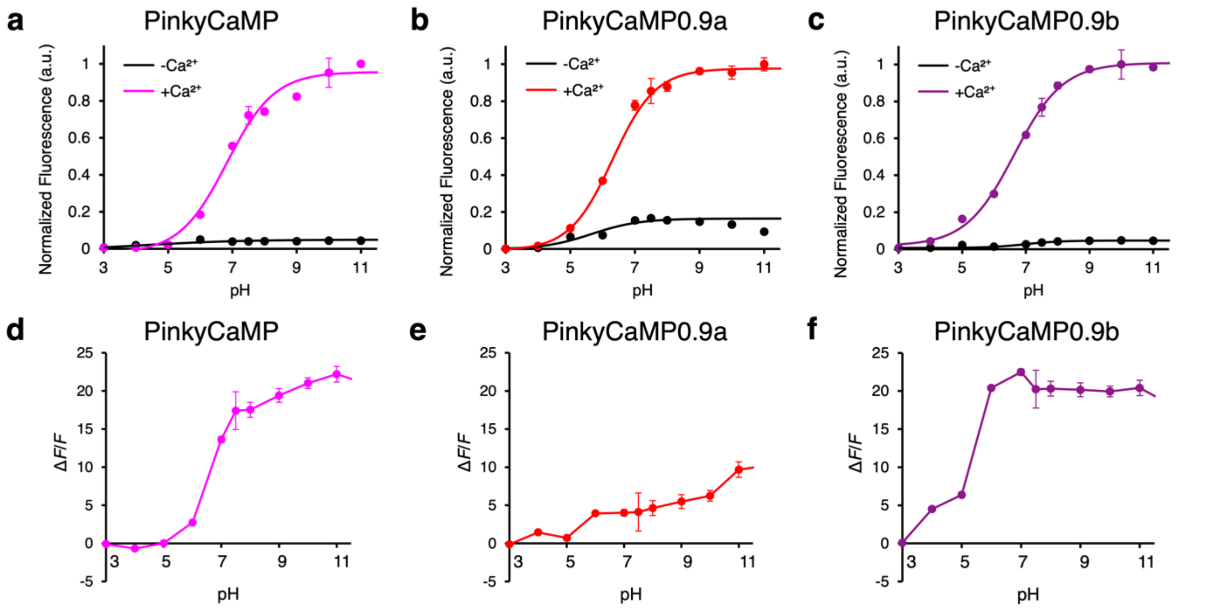


**Figure S4: pH dependence of PinkyCaMP variants.** (**a,b,c**) pH titration curves in the presence (39 µM) and absence of Ca2+ (n = 3 replicates; mean ± s.d.) for PinkyCaMP (= PinkyCaMP0.9c) (a), PinkyCaMP0.9a (b), and PinkyCaMP0.9b (c). (d,e,f) pH-dependence of the Ca2+-dependent ΔF/F for PinkyCaMP (d), PinkyCaMP0.9a (e) and PinkyCaMP0.9b (f), calculated by dividing the fluorescence intensity in the presence of Ca2+ by the fluorescence intensity in the absence of Ca2+, at a particular pH value. Lines connecting the data points are simply to guide the eye.

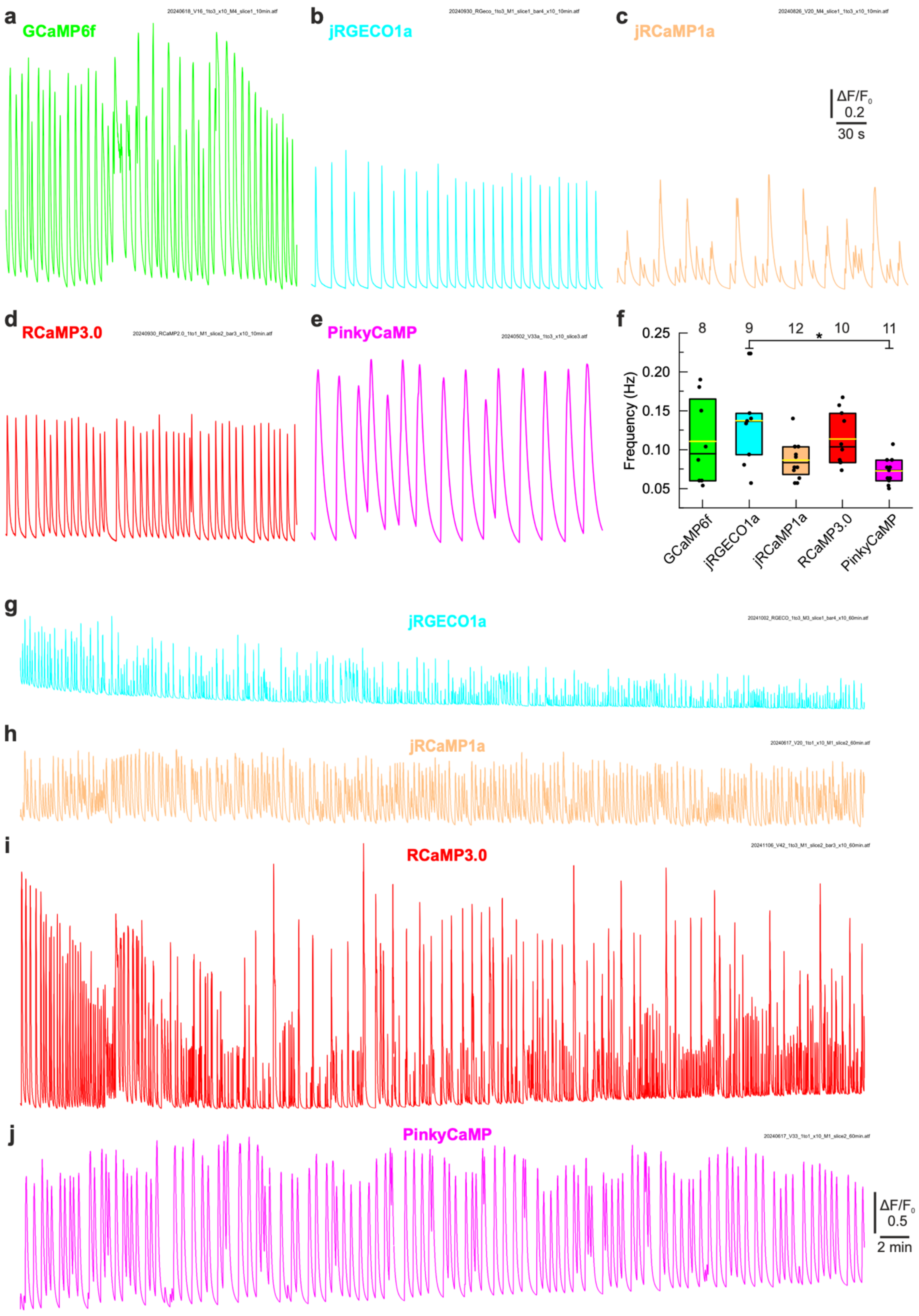


**Figure S5: Characterization of PinkyCaMP and other GECIs in organotypic slice cultures.** (a-e) Example traces (ΔF/F0) of GCaMP6f, jRGECO1a, jRCaMP1a, RCaMP3.0 and PinkyCaMP showing spontaneous synchronous network activity at DIV 13-21. The examples of RCaMP3.0 and PinkyCaMP are also shown in Fig. 3a,b. (f) Frequencies of synchronous activity (for details see Methods). The black boxes indicate 25-75% percentiles and medians, the yellow lines the means. The numbers give the numbers of analyzed slices. Statistical differences are indicated by bars with *p ≤ 0.05. (g-j) 60 min recordings of synchronous activity.

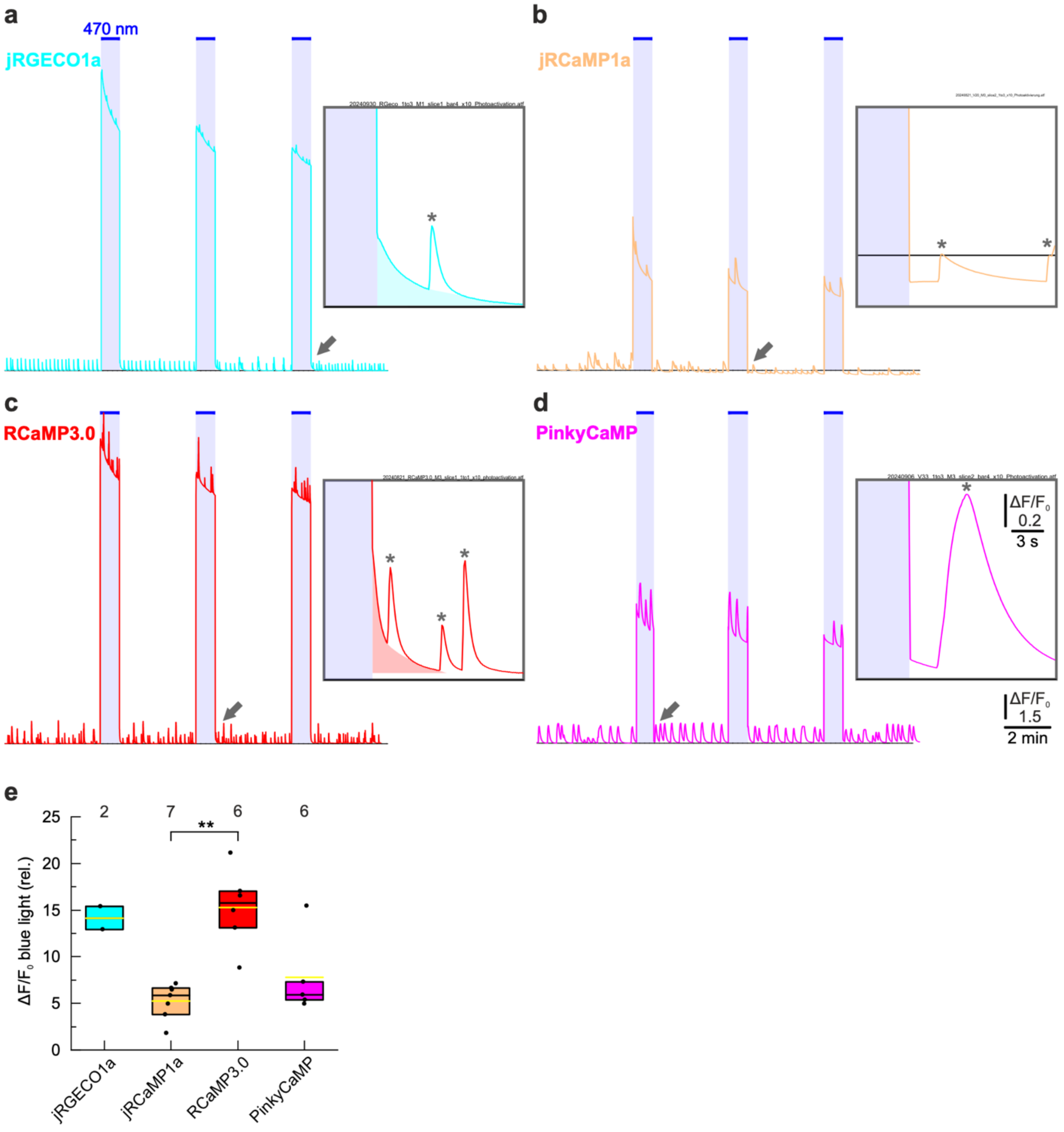


**Figure S6: Blue-light photoactivation experiments in organotypic slice cultures. (a-d)** Continuous imaging with green-light excitation (554 nm) and three additional high-intensity blue-light stimulations (470 nm, 50 s each as indicated). For details see Methods. The inserts show the first 14 seconds after the blue light is turned off. jRGECO1a and RCaMP3.0 show a decaying signal (colored background), which indicates recovery from photoactivation. These signals could be mistaken as Ca2+ elevations and have similar amplitude as synchronous events ((labeled with *). jRCaMP1a shows a slightly bleached signal, whereas PinkyCaMP shows a stable signal after the light is turned off (synchronous events again labeled with ). For quantification see Fig. 3h. (e) Fluorescence signal (emission 630 ± 38 nm) during high-intensity blue-light excitation. Blue light is more efficient in exciting fluorescence in jRGECO1a and RCaMP3.0 than in jRCaMP1a and PinkyCaMP. Black boxes indicate 25-75% percentiles and medians, the yellow lines means. The number of analyzed slices is given. Statistical differences are indicated with $^{**}p \leq 0.01$.

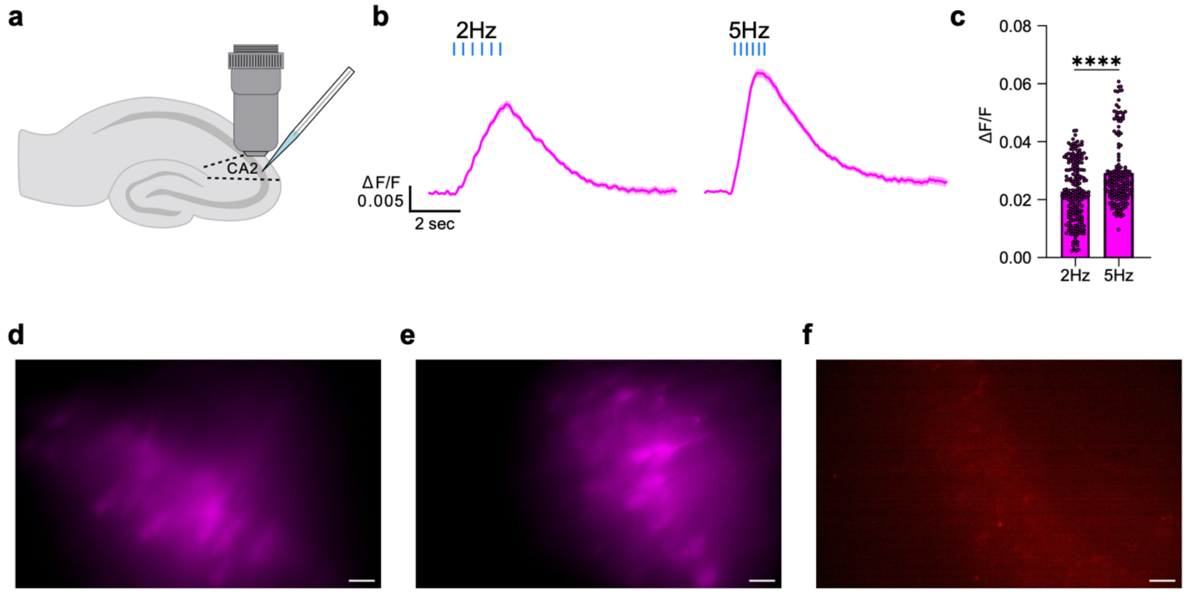


**Figure S7: Comparison of PinkyCaMP to other red GECIs in brain slices.** (a) Schematic drawing of the stimulation and recording experimental design (b) Average ΔF/F traces of 6 stimulation pulses at 2Hz (left), n=204 cells, and 5Hz (right), n=167 cells, stimulation. (c) Maximal ΔF/F values for the traces in S7(b) ($0.0228 \pm 0.0007$ and $0.0290 \pm 0.0009$ for 2Hz and 5Hz, respectively). Mann-Whitney U test, ****$p \leq 0.0001$. (d) Widefield image of PinkyCaMP in CA2 with 0.23 mW/mm² light intensity, (e) PinkyCaMP in CA2 with 11.83 mW/mm² light intensity, and (f) RCaMP3 in CA2 with 11.83 mW/mm² light intensity, scale bars 20 μm.

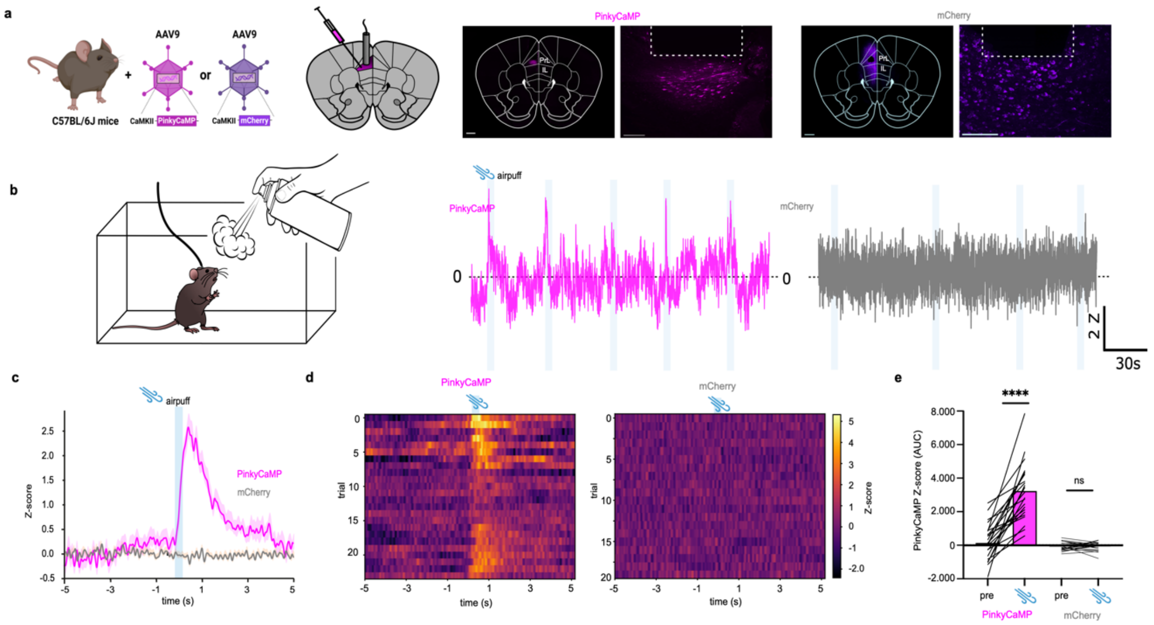


**Figure S8: Monitoring in vivo activity dynamics with PinkyCaMP.** (a) Schematic drawing of AAV injection into the prelimbic area (PrL) of the right prefrontal cortex. (b) Histology example of PinkyCaMP expression and fiber placement. Scalebar 500 µm, magnification inset scalebar 100 µm. (c) Experimental setup for airpuff application. (d) Example traces of PinkyCaMP and mCherry fluorescence in freely moving mice during an aversive airpuff in their homecage. (e) Averaged PinkyCaMP activity aligned to an aversive airpuff n=3 mice (PinkyCaMP) , n=4 mice (control) (mean ± s.e.m.). (f) Single trial heatmap of PinkyCaMP (24 trials from 3 mice) and control animals (20 trials from 4 mice). (g) Area under the curve (AUC) of PinkyCaMP and mCherry signal around the airpuff. Ordinary one-way ANOVA, PinkyCaMP: mean pre 136 ± 238; mean during: 3.246 ± 335; mCherry: mean pre 77 ± 57; mean during: -69 ± 58, **** p≤0.0001.

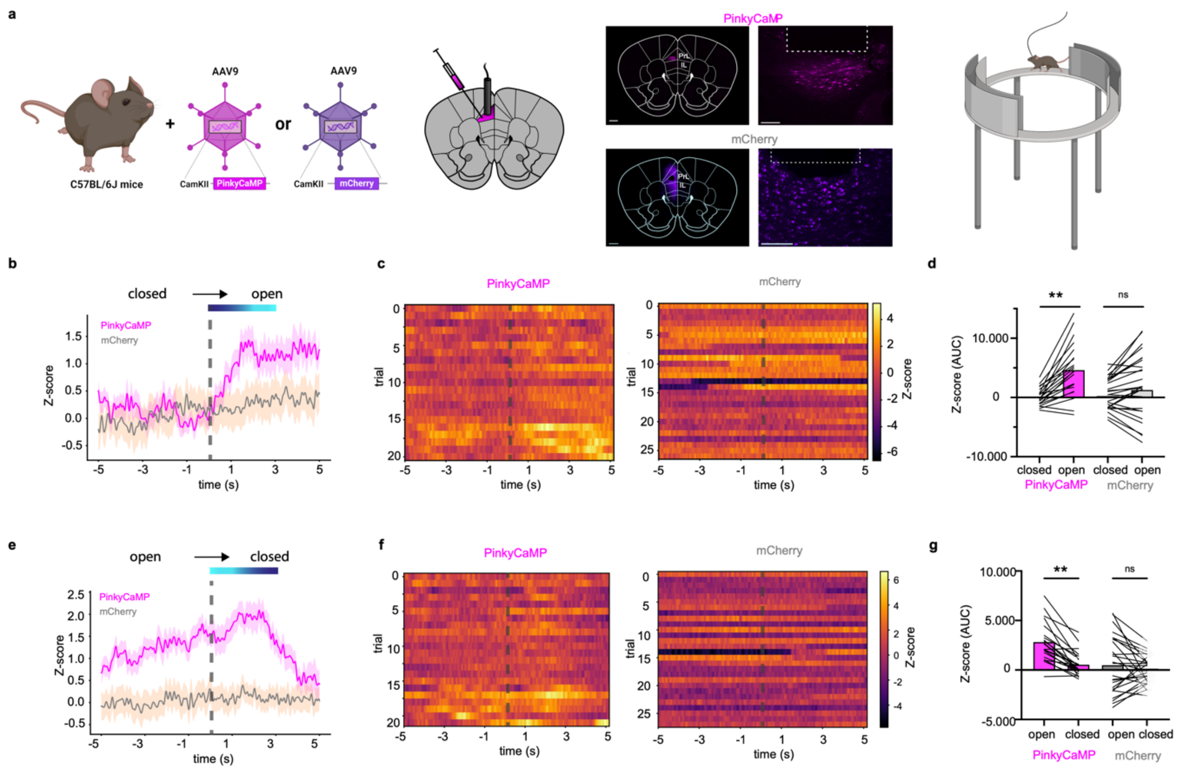

**Figure S9: Monitoring in vivo activity dynamics with PinkyCaMP in EZM.** (a) Schematic drawing of AAV injection into the prelimbic area (PrL) of the right prefrontal cortex. Histology example of PinkyCaMP expression and fiber placement. Scalebar 500 µm, magnification inset scalebar 100 µm. Experimental setup of elevated zero maze (EZM) shown on the right. (b) Averaged PinkyCaMP activity aligned to the transition from closed to open arm n=3 mice (PinkyCaMP) , n=4 mice (control) (mean ± s.e.m.). (c) Single trial heatmap of PinkyCaMP (20 trials from 3 mice) and control animals (26 trials from 4 mice) (d) Area under the curve (AUC) of PinkyCaMP and mCherry signal before and after the transition. Ordinary one-way ANOVA, PinkyCaMP: mean closed arm 90 ± 324; mean open arm: 4.708 ± 3.984; mCherry: mean closed 243± 557; mean closed: 1.336 ± 1120, ** p≤0.01. (e) Averaged PinkyCaMP activity aligned to the transition from open to closed arm n=3 mice (PinkyCaMP), n=4 mice (control) (mean ± s.e.m.). (f) Single trial heatmap of PinkyCaMP (20 trials from 3 mice) and control animals (26 trials from 4 mice). (g) Area under the curve (AUC) of PinkyCaMP and mCherry signal before and after the transition. Ordinary one-way ANOVA, PinkyCaMP: mean open arm 2.870 ± 411; mean closed arm: 576 ± 258; mCherry: mean open arm 530 ± 563; mean open arm: 136 ± 247, ** p≤0.01.

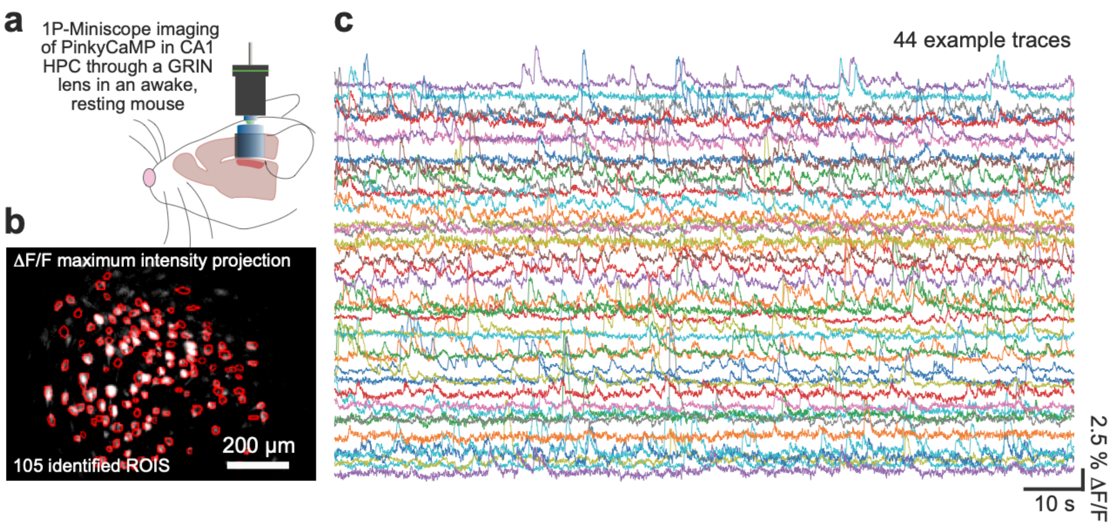


**Figure S10. Miniature microscope 1-photon imaging of PinkyCaMP expression in dorsal hippocampus.** a. Cartoon of recording setup. b. Maximum intensity projection and cell maps identified with CNMFE. c. PinkyCaMP Ca2+ fluorescence of 44 example neurons recorded in a head-fixed awake mouse with a miniature microscope for red fluorescence imaging (see Fig. 6 for two-photon imaging).
